# Gating Mechanism of β-Ketoacyl-ACP Synthases

**DOI:** 10.1101/644518

**Authors:** Jeffrey T. Mindrebo, Ashay Patel, Woojoo E. Kim, Tony D. Davis, Aochiu Chen, Thomas G. Bartholow, James J. La Clair, J. Andrew McCammon, Joseph P. Noel, Michael D. Burkart

**Affiliations:** Department of Chemistry and Biochemistry, University of California, San Diego, 9500 Gilman Drive, La Jolla CA 92093-0358; Department of Pharmacology, University of California, San Diego, 9500 Gilman Drive, La Jolla CA 92093; Jack H. Skirball Center for Chemical Biology and Proteomics, Salk Institute for Biological Studies, La Jolla, CA 92037; Howard Hughes Medical Institute, Salk Institute for Biological Studies, La Jolla, CA 92037

## Abstract

Formation of carbon-carbon bonds via β-ketoacyl-acyl carrier protein (ACP) synthases (KS), are key reactions during *de novo* fatty acid and polyketide biosynthesis. KSs recognize multiple ACPs and choreograph ping-pong mechanisms often in an iterative fashion. Therefore, KSs must limit non-productive protein-protein interactions (PPIs) to achieve high degrees of reaction fidelity. To better understand the stereochemical features governing substrate discrimination during ACP•KS PPIs, we determined x-ray crystal structures complemented by molecular dynamic simulations of *E. coli* AcpP in complex with the elongating KSs, FabF and FabB. Covalently trapped substrate analogs were used to interrogate critical catalytic events accompanying carbon-carbon bond formation revealing a previously unknown gating mechanism during the binding and delivery of acyl-AcpPs. Two active site loops undergo large conformational excursions during this dynamic gating mechanism and are likely evolutionarily conserved features generally in elongating KSs.

## Introduction

Fatty acid synthases (FASs) and polyketide synthases (PKSs) iteratively condense and reduce keto units to produce a variety of natural compounds, ranging from simple fatty acids to complex bioactive metabolites.^1,2^ FASs and PKSs can exist as one or more polypeptide “mega-synthases” that contain distinct catalytic domains (type I) or as discrete enzymes, each possessing a single activity (type II).^3^ In FASs and PKSs, β-ketoacyl-acyl carrier protein synthases, alternatively ketosynthases (KSs), catalyze carbon-carbon formation via an efficient and controlled decarboxylative Claisen-like condensation reaction. The formation of carbon-carbon bonds not only leads to the production of complex natural products but also provides a thermodynamic driving force for their formation.^4,5^

Elaboration of these natural products is facilitated by a family of small ~8 kDa proteins, acyl carrier proteins (ACPs), that shuttle fatty acid and polyketide substrates and intermediates as they are processed by catalytic domains or discrete enzymes.^6,7^ ACPs are translated in inactive *apo* forms and are posttranslationally modified to active *holo* forms with a 4’-phosphopantetheine (PPant) arm attached at a conserved serine residue, providing a thiol moiety that ligates substrates and intermediates to the ACPs.^6^ Protein-protein interactions (PPIs) between ACPs and their enzymatic partners (or domains) regulate the catalytic activities of FASs and PKSs kinetically and stereochemically.^3,8–13^ Therefore, it is important to elucidate the structural and biophysical underpinnings of the PPIs between ACPs and their cognate KSs to understand reaction trajectories and product fidelity in FAS and PKS biosynthetic pathways.

In contrast to initiating KSs^14,15^, which prime *holo-*ACPs, elongating KSs, can iteratively process fatty acid or polyketide intermediates.^3,16^ Despite their distinct biosynthetic roles, these KSs operate via a common reaction mechanism, which can be separated into two half-reactions.^17,18^ In the first half-reaction, an acyl-ACP associates with a KS, and its acyl cargo is transferred from the prosthetic PPant arm to a conserved, active site cysteine, producing a covalent acyl-KS intermediate and *holo*-ACP. Malonyl-ACP then replaces *holo*-ACP and undergoes a decarboxylative Claisen-like condensation with the acyl-KS adduct, producing and offloading β-ketoacyl-ACP^17^ (Figure S1). During the course of this ping-pong process, two distinct ACPs must sequentially interact with the KS enforcing exquisite temporal and spatial discrimination among subtly different chemical states of these acyl-ACPs.

The *Escherichia coli* type II FAS system has served as a model for understanding ACP-mediated PPIs as well as the enzymatic transformations catalyzed by FASs and PKSs (Figure 1).^19–23^ Herein, we utilize *E. coli* fatty acid biosynthesis (FAB) to further elucidate the mechanisms of elongating KSs. AcpP, *E. coli*’s FAB ACP, is the most abundant cellular protein, interacts with at least 21 enzymatic partners^24,25^ (Figure S1), and yet, must facilitate the production of enough fatty acids to support *E. coli* doubling-times of 20 minutes.^26^ Therefore, AcpP and its partner enzymes must minimize the number of non-productive PPIs. The KS, in particular, must precisely coordinate ACP binding events to catalyze carbon-carbon bond formation efficiently while also controlling product chain lengths.^27–29^ We used this well-defined system to better understand ACP•KS PPIs and KS substrate discrimination by structurally characterizing *E. coli* elongating KSs as substrate/intermediate complexes that approximate states formed during fatty acid elongation.

**Figure 1.**
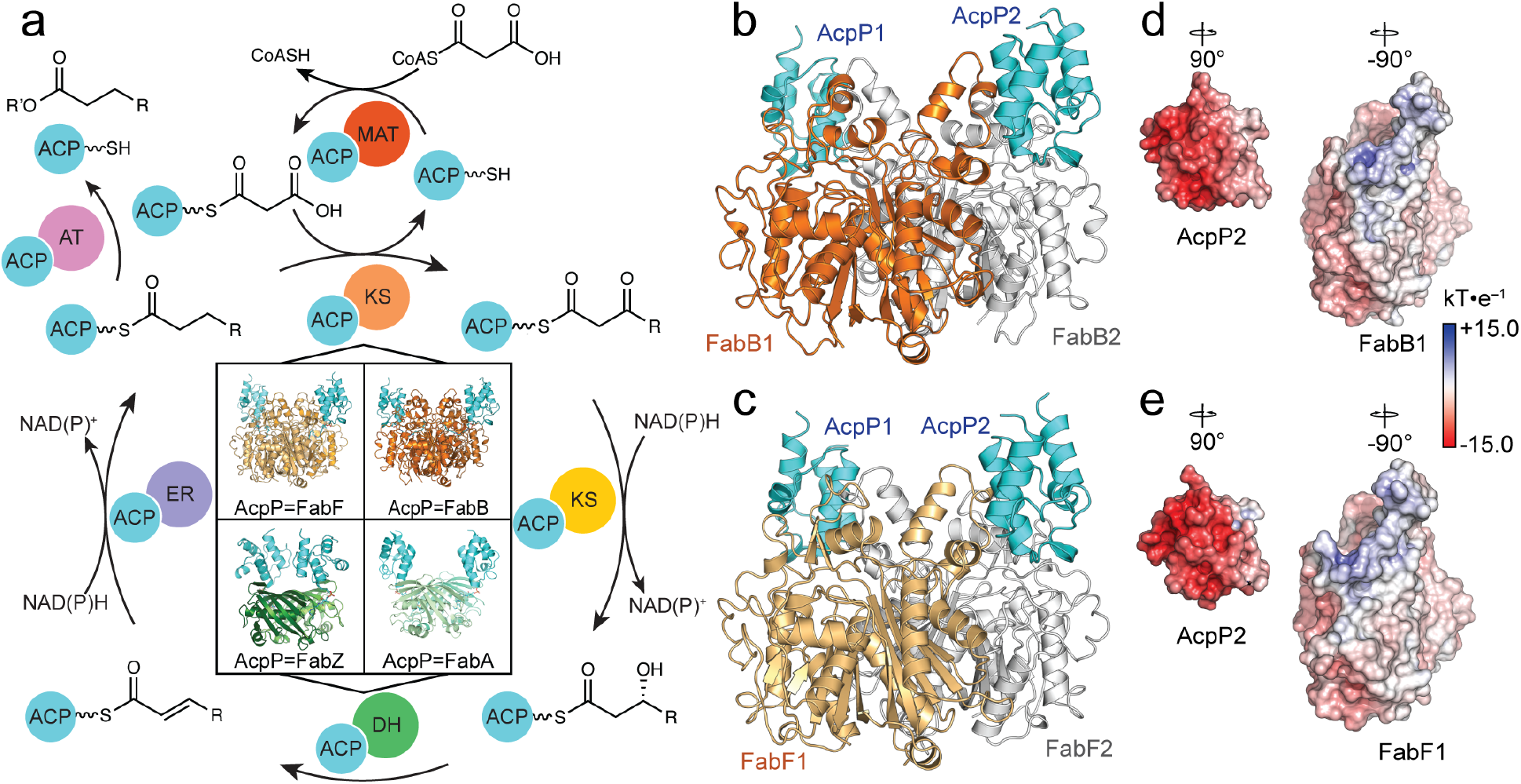
FAB cycle and known AcpP=PP complexes. **a**, A schematic representation of the chemical transformations that occur during fatty acid biosynthesis, highlighting the importance of protein-protein interactions required at each step for substrate processing. The four quadrants at the center of the scheme show the structures of all FAS AcpP-PP complexes solved thus far. The top two structures are the crosslinked AcpP-KSs (AcpP-FabF and AcpP-FabB) reported herein, while depicted below are the crosslinked AcpP-DHs (AcpP-FabZ and AcpP-FabA) solved in previous publications. ACP = acyl carrier protein, MAT = malonyl-ACP transacylase, KS = ketosynthase, KR = ketoreductase, DH = dehydratase, ER = enoylreductase, AT = acyl transferase. AcpP-FabZ (PDB: 6N3P), AcpP-FabA (PDB: 4KEH). ***b,c***, Cartoon representations of the crosslinked AcpP-FabB and AcpP-FabF dimers. AcpP is colored cyan; the first KS monomer of AcpP-FabB and AcpPFabF is colored orange or light orange, respectively, while the second is colored white in both structures. ***d***, Electrostatic potentials (ESP) maps of the FabB and AcpP monomers. ***e***, ESP map of the FabF and AcpP monomers. In all cases, the ESP is mapped onto the “Connelly” surfaces of the FabF and AcpP monomers using a blue to white to red color range, spanning from +15.0 kT•*e*^−1^ to –15.0 kT•*e*^−1^.

Here we report newly described stereochemical features of the mechanisms of two *E. coli* elongating KSs, FabB and FabF, representing the KASI and KASII KS families, respectively. Site-specific crosslinking probes were employed that mimic key catalytic intermediates to facilitate elucidation of the structures of AcpP•KS complexes by protein x-ray crystallography. In conjunction with molecular dynamics (MD) simulations, these structures provide the first dynamic descriptions of two ACP•KS complexes revealing conformational changes that regulate substrate processing. This mechanism employs two conserved active site loops that open and close through a double drawbridge-like gating mechanism.^30^ When open, the drawbridge expands the KS active site in order to accommodate large PPant-tethered acyl-AcpP substrates. The structural features underlying this gating machinery are conserved in related enzymes; therefore, similar conformational transitions will likely determine substrate selectivity and processing in other FAS and PKS biosynthetic pathways.

## Results

### Crosslinking, Crystallization, and MD simulations of AcpP•KS Complexes

We synthesized and chemoenzymatically loaded C8-α-bromo-, C12-α-bromo-, and C16-α-bromopantetheineamide crosslinking probes onto AcpP to produce C8αBr-*crypto*-AcpP (C8AcpP), C12αBr-*crypto*-AcpP (C12AcpP), and C16αBr-*crypto*-AcpP (C16AcpP) (Figure S1).^31–33^ These *crypto*-AcpPs were crosslinked with FabF and FabB to produce crosslinked complexes (AcpP-KS), which were then crystallized and analyzed at the Advanced Light Source (ALS). C12AcpP-FabF and C16AcpP-FabF samples provided 2.35 Å and 2.30 Å resolution data sets, respectively. The crystallographic asymmetric unit (ASU) of these complexes were identical and contained one monomer of FabF crosslinked to a single AcpP. Additionally, two different C12AcpP-FabB crystals diffracted to 1.55 Å and 2.20 Å, respectively, while C16AcpP-FabB diffracted to 2.50 Å. All of the AcpP-FabB complexes crystallized in different space groups (Table S1). The 1.55 Å C12AcpP-FabB and 2.50 Å C16AcpP-FabB ASUs contain functional FabB dimers crosslinked to two AcpP molecules while the 2.2 Å C12AcpP-FabB crystal contains two functional FabB dimers and four crosslinked AcpPs per ASU.

Using these AcpP-KS crystal structures as initial coordinates, we next performed MD simulations of 10:0, 12:0, 16:0-AcpP•KS complexes (non-crosslinked) (Figure S1) to probe the dynamic properties accompanying the PPIs between the acyl-AcpPs and KSs. Models of the noncovalent acyl-AcpP-KSs were constructed by manually replacing the crosslinked chemical probe present in the crystal structures with the mimicked acyl substrate (Online Methods). In addition, simulations of *apo*-FabB and *apo*-FabF were performed using previously reported crystal structures (PDB: 2V69 and 2GFW, respectively) as initial coordinates. Each system was subjected to three independent 500 ns MD simulations. A total of 12 μs of MD data was collected.

### The AcpP-KS Interface

The AcpP-KS interacting surfaces are electrostatic in nature (Figure 1b-e) composed largely off flanking electrostatic interactions and hydrogen bonds surrounding a central hydrophobic patch as shown in Figures 2 and S2-S4. The AcpP and KS monomers provide negatively and positively charged residues, respectively, to the interface. Milligan *et al.* have recently reported a structure which delineates the AcpP-FabB interface (PDB: 5KOF). Herein, we report the first example of a KASII-type KS in complex with an ACP, AcpP-FabF. The AcpP-FabF complex shows one region (region 1) encompassing Lys65, Arg68, and Lys69 of FabF interacting with Glu13, Gln14, Asp35, and Asp38 of AcpP. Additionally, Thr270 of FabF, which sits on a flexible loop that undergoes conformational rearrangements upon acyl-AcpP binding, interacts with Asp35 and Ser36 of AcpP. A second region (region 2) comprising Arg127, Lys128, and Arg206 of FabF contacts Glu47, Glu48, Glu53, and Asp56 of AcpP. The central hydrophobic patch is comprised of interactions between Met44, Val40, and Leu37 of AcpP and Ile129, Ser130, Pro131, Phe132, Ala205, and Arg206 of FabF (Figure S2e,h).

**Figure 2.**
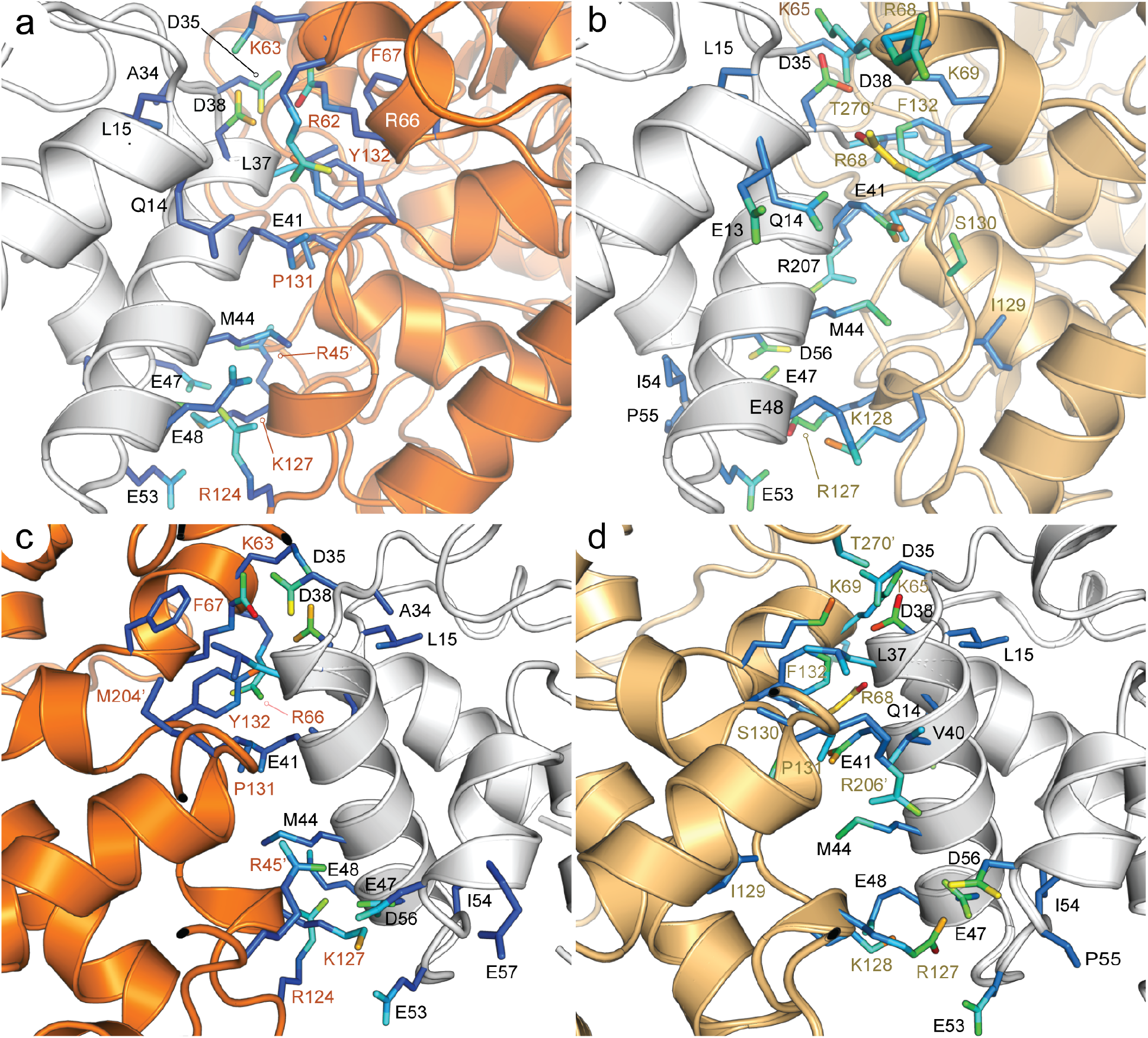
The structure and interface of AcpP=FabB and AcpP=FabF. **a**, A close-up view of the interface between AcpP-FabB. ***b***, A close-up view of the interface between AcpP and FabF. ***c***, An alternate view of the interface between AcpP-FabB, related to the view shown in panel ***a*** by 180° rotation about the vertical. ***d**,* An alternate view related to the view shown in panel ***b*** by 180° rotation about the vertical of the interface between AcpP and FabF. In panels ***a***-***d***, interfacial residues (determined computationally) are shown as sticks; residues of AcpP and FabB (or FabF) that “engage” in intermolecular contact(s) with the partner protein are labeled in black and dark orange (or light orange), respectively. The color of the side chain atoms of interfacial residues indicates the frequency at which the atom is sampled computationally as forming an intermolecular contact. This color ranges from purple (rarely in contact) to red (often in contact).

Comparisons of the AcpP-FabF and AcpP-FabB PPI interfaces reveal that they share similar chemical features, but structural alignments of the KS subunits show that the orientations of AcpPs in FabF and FabB complexes are distinct (Figure S2a,b). In FabF, AcpP1 and AcpP2 bury surface areas of 671.2 Å^2^ and 675.4 Å^2^, respectively, while in FabB, AcpP1 and AcpP2 bury surface areas of 597.5 Å^2^ and 607.4 Å^2^, respectively. These small contact areas are consistent with the transient nature of ACP interactions^10,23^. In addition, the differences in contact areas of AcpP-FabF and AcpP-FabB interfaces are consistent with the differing positions that AcpPs adopt relative to their KS partners (Figure S2a,b).

MD simulations of the various acyl-AcpP-KS complexes (Figure 2a-d) show the extent to which atoms interact intermolecularly at the AcpP-KS interface. FabF, unlike FabB, forms long-lived contacts with AcpP’s helix I residues Glu13 and Gln14 during the course of the simulations (Figure 2a-d, Figures S3,S4). FabF forms additional short-lived hydrophobic contacts with AcpP throughout the MD simulations, consistent with the larger contact area of the AcpP-FabF interface. (Figures S3,S4) Simulations also reveal that the charged residues of regions 1 and 2 of the AcpP-KS interface form and break interactions with one another throughout the course of the simulation (Figure 2a-d and Figures S2-S4), showcasing the dynamism of the PPIs of these complexes. Simulations show that AcpP’s Asp35 and Asp38 are the residues most frequently forming interactions at the AcpP-FabB interface (Figure S3, S4).

### Conformational heterogeneity in FabF active site loops

We could not definitively locate density for the alkyl chain of the crosslinking probe in the C12AcpP-FabF structure. This lack of density is accompanied by substantial conformational heterogeneity in the nearby GFGG β-turn (residues 399-402), referred to herein as loop 1, and a second loop (residues 265-275) buttressed by loop 1, referred to herein as loop 2. Therefore, we turned to our 2.30 Å C16AcpP-FabF dataset, which provided electron density maps of sufficient quality to model and refine the C16-acyl mimetic and the previously unresolved loops 1 and 2 (Figures 3,4, S5a-c). In C16AcpP-FabF, both loops move to create a hydrophobic pocket that accommodates the PPant-tethered acyl group (Figure 4a,f, Figure S5a-c). The C16-acyl chain electron density delineates two conformations (Figure 3b, Figure S5a-c), neither of which place the substrate in the acyl binding pocket as seen in the previously reported dodecanoyl-FabF (PDB: 2GFY) structure (Figure 3c).^34^ The carbonyl group of the fatty acyl chain resides within hydrogen-bonding distance of the two catalytic histidine residues of FabF (His303, His340) (Figure 3b), suggesting the C16AcpP-FabF structure resembles an intermediate state formed before transacylation or after condensation with malonyl-AcpP.

**Figure 3.**
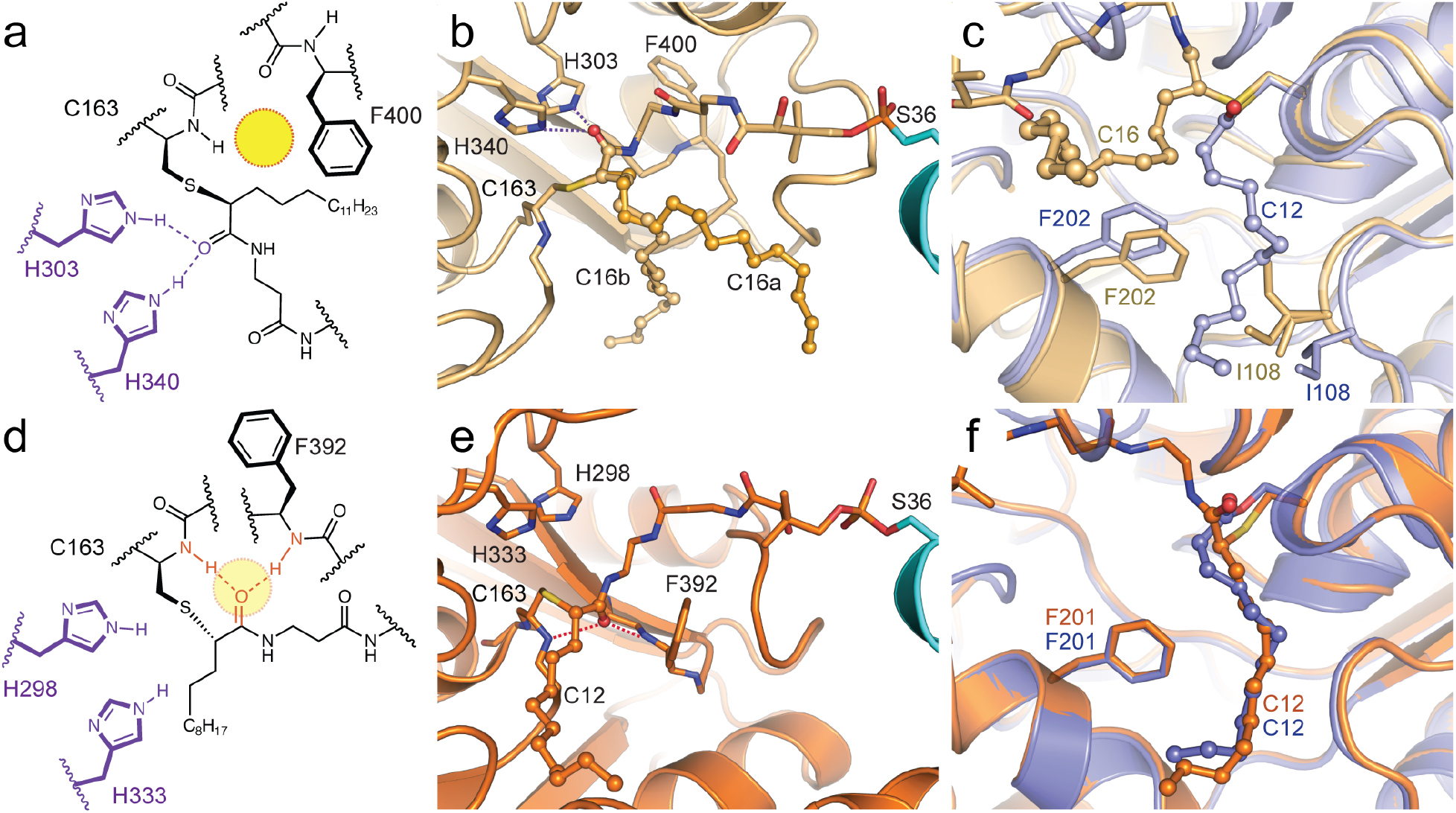
FabF active site interactions with C16 substrate analog. ***a***, 2D representation of the 4’-phosphopantetheineamide crosslinking probe within the FabF active site. ***b***, 3D rendering of the C16AcpP-FabF crystal structure illustrating the two distinct C16-acyl binding modes, C16a (ball and stick) and C16b (ball and stick), assumed by the aliphatic portion of the probe. AcpP and FabF are colored cyan and light orange, respectively; the 4’-phosphopantetheinyl arm of the probe is represented as sticks, while the acyl chains are represented in ball and stick form. Active site catalytic residues, Cys163, His303, His340, and Phe400, are represented as sticks and colored according to element. ***c***, Comparison of acyl binding pocket and associated gating residues, F202 and I108, in dodecanoyl-FabF (12:0-FabF) (2GFY, light purple), and C16AcpP-FabF (light orange). ***d***, 2D representation of the 4’-phosphopantetheine crosslinking probe within the FabB active site. ***e***, 3D rendering of the C12AcpP-FabB crystal structure illustrating the probe an acyl chain binding mode. ***f***, Comparison of acyl binding pocket and in dodecanoyl-FabB (12:0-FabB) (PDB ID: 1EK4, bright purple), and C12AcpP-FabB (bright orange). Residues in panels are numbered according to their respective protein, FabF is numbered in FabF residue numbering and FabB is numbered in FabB residue numbering. Hydrogen-bonding interactions are represented as dotted lines.

**Figure 4.**
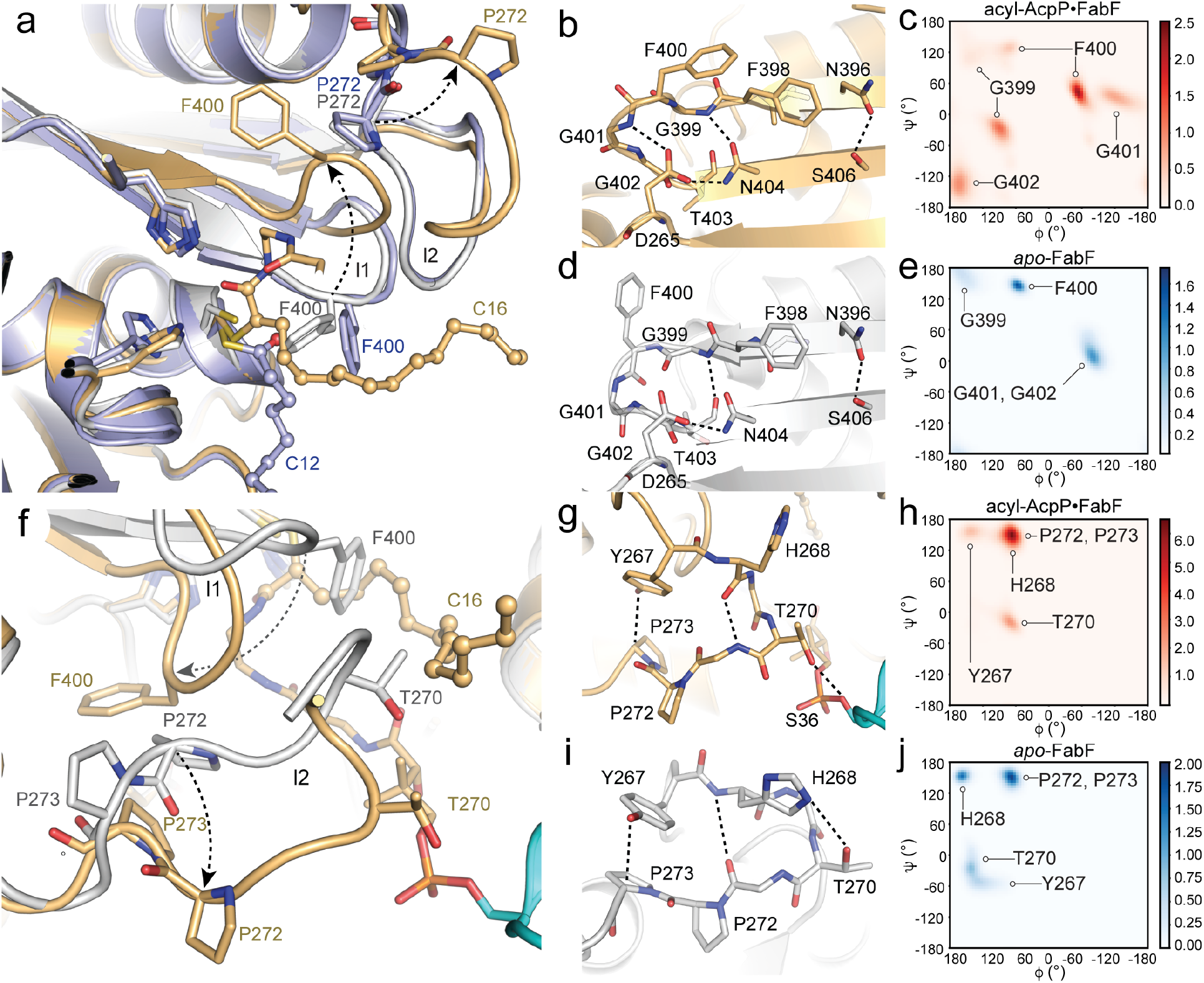
Conformational changes in FabF active site loops. ***a***, Comparison of F400, loop 1, and loop 2 conformations of *apo*-FabF (light grey, PDB: 2GFW), dodecanoyl-FabF (light purple, PDB: 2GFY,), and C16AcpP-FabF (light orange). ***b***, Interactions that stabilize the open-conformation of the β-turn motif of loop 1 in C16AcpP-FabF (light orange). ***c***, Ramachandran analysis of the conserved GFGG sequence within loop 1 from 1.5 μs of 10:0-AcpP•FabF, 12:0-AcpP•FabF, 16:0-AcpP•FabF using 4.5 μs of simulations data. ***d***, Interactions that stabilize the closed-conformation of the β-turn motif of loop 1 in *apo*-FabF (light grey). In **a**, **b**, **d**, **f**, **g**, and **h**, loop 1 residues are shown as sticks and colored according to element. Hydrogen-bond interactions are highlighted using dotted lines. ***e***, Ramachandran analysis of the conserved GFGG sequence within loop 1 from 1.5 μs of MD simulation data of *apo*-FabF. In ***c*** and ***e***, simulated backbone ψ and φ dihedral were binned using widths of 5°. Color bars indicates density of data points within each 2D bin. ***f***, Loop 2 overlay of *apo*-FabF (light grey, PDB: 2GFW) and C16AcpP-FabF (light orange) with the crosslinked AcpP colored cyan. ***g***, Interactions that stabilize the β-turn “open conformation” of loop 2 in C16AcpP-FabF (light orange). ***h***, Ramachandran analysis of key loop 2 residues of FabF monomers of acyl-AcpP•FabF complexes (“gate-open” conformation). ***i***, Interactions that stabilize the loop 2 “closed conformation” in *apo*-FabF (white). **j**, Ramachandran analysis of key loop 2 residues of FabF of *apo*-FabF (“gate-closed” conformation).

Unlike C16AcpP-FabF, MD simulations suggest that the acyl chain carbonyls in 10:0-AcpP-FabF, 12:0-AcpP-FabF, 16:0-AcpP-FabF complexes do not form hydrogen bonds with either active site histidine (Figure 3a-c). Over the course of the simulation, the substrate relaxes to assume a distinct (pre-transacylation/post-condensation) disposition within the hydrophobic pocket described above (Figure S5). Furthermore, the oxyanion hole comprising Cys163 and Phe400 backbone amides was not engaged by the substrate during the course of the simulations (Figure S6). These findings, in addition to long distances (d > 5.0 Å) between the substrates and the thiolate moiety of the active site cysteine (Figure S6), suggest the AcpP-FabF structure features a pose wherein the crosslinker mimics a substrate conformation prior to transacylation, or a production conformation before dissociation.

### FabF active site loops adopt a “gate-open” conformation to accommodate acyl cargo

Comparison of the *apo*-FabF (PDB: 2GFW), acyl-FabF (PDB: 2GFY), and C16AcpP-FabF structures shows that Phe400 moves ~7 Å, as measured by changes in Cα positions, into a pocket created by the displacement of loop 2 (Figure 4a,f). Gly399 and Gly402 of loop 1 act as pivot points as the loop transits from a closed conformation to an open conformation. (Figure 4b-e). The open conformation is stabilized by a network of hydrogen-bonding interactions between the backbone amides of Gly399, Phe400, Gly401, and Gly402 and the side chains of Asp265 and Asn404 (Figure 4b,d). Overlays of the *apo*-FabF and C16AcpP-FabF structures indicate that the two observed conformations of the C16-acyl chain clash with Phe400 in the closed loop 1 conformation (Figure 4a).

Loop 2 pivots about a Pro-Pro motif, whereby Pro273 undergoes a 165° change in its ψ angle, causing the loop to swing outward by 5 Å (Figure 4f). Analysis of loop 2 backbone dihedrals suggests the loop transitions from a type VIII β-turn in the closed conformation to a type I β-turn in the open conformation. In the closed conformation, loop 2 is stabilized by a Ser271_(O)C_-His268_(H)N_ main-chain hydrogen bond, a Thr270-His268 side-chain hydrogen bond, and a Pro273-Tyr267 C-H π interaction^35^ (Figure 4g,i). Transition to the open conformation causes the β-turn backbone hydrogen-bonding pattern to flip from Ser271_(O)C_-His268_(H)N_ to Ser271_(N)H_-His268_(O)C_ and Thr270 breaks its hydrogen bond with His268 to interact with Asp35 and the γ-hydroxyl of Ser36 from AcpP.

The coordinated movement of loops 1 and 2 of FabF resembles a double-drawbridge gate that regulates active site access for acyl-AcpPs (Figure 4a,f).^30^ The residues that define this pocket are hydrophobic and serve to stabilize the aliphatic cargo of AcpP as it is “de-sequestered” or “re-sequestered” during chain-flipping^25^ (Figure S5b,c). Furthermore, loop 1 only forms the active site oxyanion hole when in the gate-closed formation, which indicates that the gating machinery not only controls ligand access to the active site but also elements of catalysis.

### Acyl-AcpP-FabB locks into a catalytically competent, “gate-closed”, acyl transfer conformation

Analysis of the 1.55 Å C12AcpP-FabB structure demonstrates well-defined electron density for the entire 4’-phosphopantetheineamide arm, the C12-acyl chain of the probe (Figure S5d), and loops 1 and 2 which exist in a catalytically competent, gate-closed conformation (Figure 3d,e). Cys163 is covalently attached to the α carbon of the acyl chain and the probe’s carbonyl group resides in the oxyanion hole formed by the backbone amides of Cys163 and Phe400 of loop 1 (Figure 3d,e). Note, the recently deposited AcpP-FabB structure (PDB: 5KOF) does not feature these interactions, as the C3-chloroacrylate probe used to trap that complex forms hydrogen bonds with His298 and His333, as seen in the AcpP-FabF structures reported herein. The acyl chain accesses the acyl binding pocket through a gate created by Phe201 and a flexible glycine-rich loop (residues 106-108) (Figure 3f, S5e), similar to the previously published dodecanoyl-FabB structure (PDB: 1EK4).^21^ Only the C12AcpP-FabB structure reported herein illustrates a ACP-KS complex in a state preorganized to catalyze transacylation.

The 2.50 Å C16AcpP-FabB structure reveals a lack of electron density for the C16-acyl chain in the acyl binding pocket, despite showing interpretable electron density for the 4’-phosphopantetheineamide portion of the probe (Figure S5f). A C12-acyl chain was modeled into the acyl binding pocket, but low levels of electron density were only observed for the alkyl chain in 2Fo-Fc maps at or below contour levels of 1σ (Figure S5f). Moreover, extension of the modeled C12 substrate to C16 reveals a constricted fatty acid binding pocket that places the alkyl chains within either KS monomer in close proximity to one another and in unfavorable conformations (Figure S5g). Additionally, clashes between the terminal carbons of the C16-acyl chain with Glu200 and Gln113 along the back wall of the binding pocket restrict accommodation of the C16 substrate. Interestingly, in our C12AcpP-FabB datasets, the end of the C12 chain stops just short of these residues (Figure S5e), indicating that they may play a role in the substrate specificity of FabB.^36–39^

MD simulations of the acyl-AcpP-FabB complexes suggest that the dynamic features of these active sites are consistent with an active site preorganized for transacylation. The active site cysteine’s thiolate moiety is significantly closer to the electrophilic carbon of the substrate (ca.3.0 Å < d < 5.5 Å) than is observed in MD simulations of acyl-AcpP-FabF complexes (Figure S6). Backbone hydrogen-bonding interactions involving the substrate, Cys163, and Phe400 were only maintained in simulations of 12:0-AcpP-FabB (Figure S7). These interactions are disrupted during simulations of 16:0-AcpP-FabB, whereas in simulations of 10:0-AcpP-FabB, the substrate is ~5 Å away from either backbone hydrogen-bonding donor. Finally, no hydrogen bonds between the active site histidines (His303 and His340) and any substrate modeled are sampled during MD simulations (Figure S7).

### Conservation of loop 1 and loop 2 residues highlights their mechanistic importance in KS catalytic activity

Multiple sequence alignments (MSA) of FabFs and FabBs indicate loop 1 is highly conserved while loop 2 is only conserved within, but not between, the two families of condensing enzymes (Figure S8a). The consensus sequence for loop 1 is comprised of the conserved GFGG β-turn motif flanked by two equally conserved asparagine residues, Asn396 and Asn404 (FabF numbering). Asn404 interacts with Asp265 found at the start of loop 2 in both the open and closed states, linking the movement of loops 1 and 2. In the open conformation, Asp265 and Asn404 form a hydrogen-bonding network with the backbone amides of Gly399, Gly401, and Gly402 in the loop 1 β-turn (Figure 4d). The highly conserved nature of loop 1 suggests that the proposed gating mechanism is a general feature of elongating KSs.

MSA analysis of FabF orthologues shows modest loop 2 conservation with the consensus sequence being D-A-Y-H-I/M-T-A-P-X-X-X-G where residues Asp265, His268, and Thr270 are highly conserved and an aromatic residue, usually tyrosine, is always found at position 267 (Figure S8b). Loop 2 from the FabB orthologues is two amino acids shorter than loop 2 of FabF and the only conserved residues between the two KS families are Asp265 and Pro272. Therefore, the less conserved loop 2 may serve as the regulatory loop during gating events as it caps the more conserved loop 1 region and could potentially modulate the rate of gate sampling or assist in the selection of specific substrates in different KSs.

### MD simulations support a gating mechanism involving loops 1 and 2

Ramachandran analyses of the GFGG motif’s fluctuations during MD simulations illustrate the rigidity of loop 1 in both the closed (*apo*-FabF, *apo*-FabB, and acyl-AcpP-FabB) and open (acyl-AcpP-FabF) conformations. The ψ and φ angles of the GFGG residues sampled computationally are narrowly distributed (Figure 4c,e) with hinge residues, Gly399 and Gly402, possessing distinct backbone dihedral angles in the open and closed conformations. MD simulations show that although the ψ and φ angles of Pro272 and Pro273 of loop 2 are distinct in the open and closed states in our crosslinked structures, they sample similar ψ, φ angle distributions (Figure 4h,j).

In an attempt to visualize the transition between the open and closed conformations of loops 1 and 2, we performed additional MD simulations of *apo*-FabF and *apo*-FabB using KS structures prepared by removing the AcpP monomers from the AcpP-KS complexes herein. We distinguish these structures from the crystallographic *apo*-FabB (PDB: 2V69) and *apo*-FabF (PDB: 2FGW) structures by referring to them as *apo*-FabB* and *apo*-FabF*. Because these *apo-* structures feature their loops in conformations that may not be stable in the absence of an associated ACP, we hypothesized that they may more readily undergo conformational transitions.

Unfortunately, no such transition was sampled during the course of any of the 3 independent 500 ns simulations performed using either KS, suggesting that the correlated movement of loops 1 and 2 between their open and closed states is a relatively slow process, occurring on timescales of 10’s or 100’s of μs.^40^ Nonetheless, analysis of the per residue backbone and side chain root mean square fluctuations (RMSFs) of the loop residues, shown in Figure 5d-i, suggests the following. Firstly, while the loops of *apo*-FabB and *apo*-FabF have similar RMSFs, the loops of *apo*-FabF* and the acyl-AcpP-FabF complexes show greater fluctuations than do those of *apo*-FabB* and the acyl-AcpP•FabB complexes, respectively. Of the structures shown in Figure 5d-i, *apo*-FabF* is the most interesting. RMSF analysis shows that loop 2 of *apo*-FabF* fluctuates more in this structure than any other simulated structures. Despite this dynamism, *apo*-FabF*’s loop 1 remains fairly rigid. It is possible that the transition of loop 2 from its open to closed conformation first requires the motion of loop 1 toward the active site. In fact, this movement of loop 1 would restore the oxyanion hole absent in the AcpP-FabF structure. Thus, the large RMSFs observed for *apo*-FabF*s loop 2 reveal a “frustrated” loop unable to assume its “preferred” open conformation in the absence of acyl-AcpP (Figure 5). These findings along with published studies of PKS ACPs^12^ suggest that ACPs may not only be responsible for substrate delivery but may also induce loop motions supporting catalysis through allostery.

**Figure 5.**
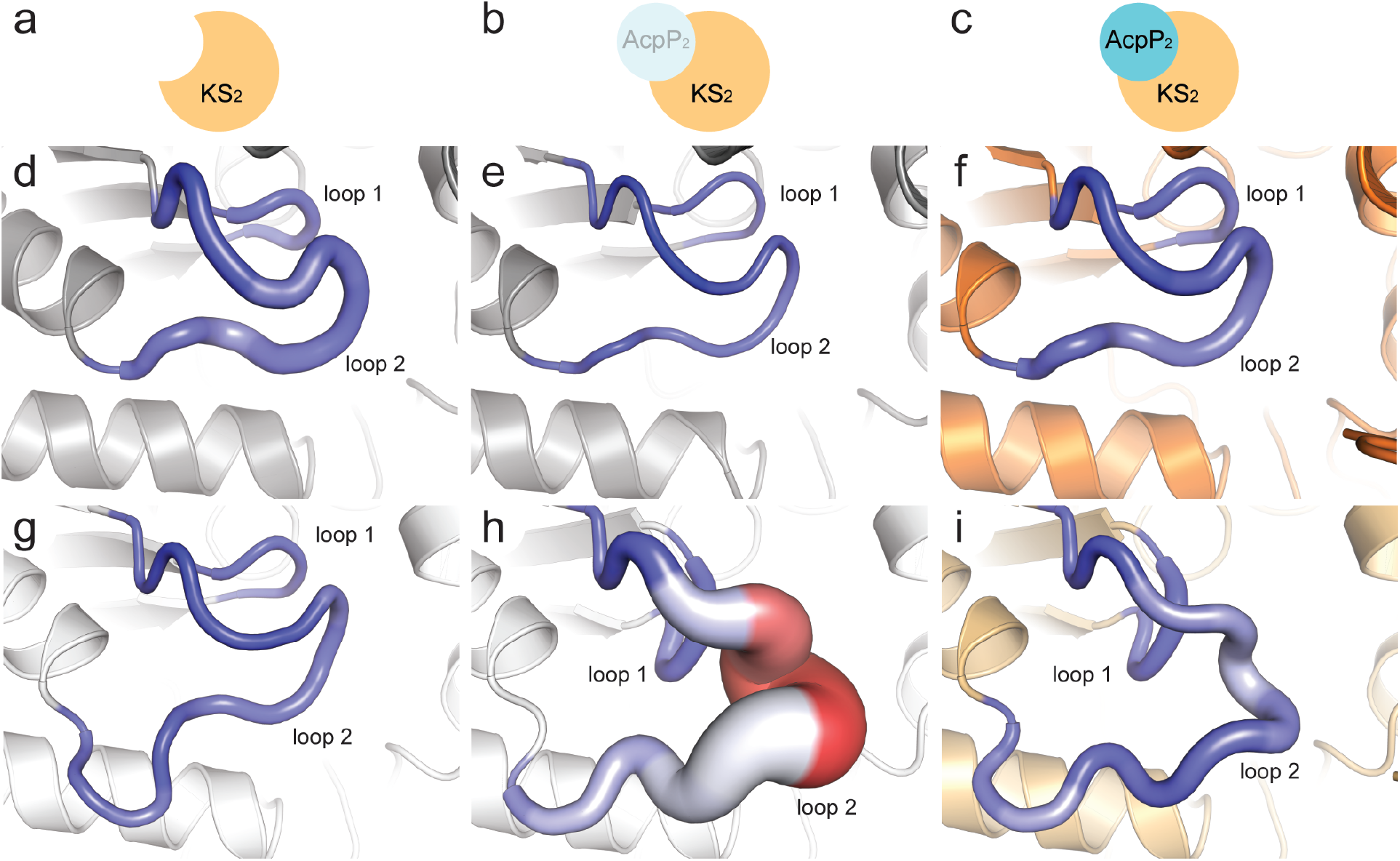
Loop dynamism sampled via computer simulations. **a**, *apo*-KS2. **b**, *apo*-KS2* (*apo* structures derived from AcpP_2_-KS_2_ structures). **c**, crosslinked AcpP _2_-KS_2_. ***d***, Root mean square fluctuations (RMSFs) of *apo*-FabB2 (determined from simulations) “mapped” onto loops 1 and 2 from the *apo*-FabB2 structure (PDB: 2GFW). ***e***, RMSFs – determined from simulations – of *apo*-FabB2* (apo structure derived from AcpP_2_-FabB_2_) mapped onto loops 1 and 2 from the *apo*-FabB2 structure. ***f***, RMSFs determined from simulations of acyl-AcpP_2_•FabB_2_ mapped onto the crystal structure of crosslinked AcpP_2_-FabB_2_. ***g***, RMSFs of *apo*-FabF2 (determined from simulations) “mapped” onto loops 1 and 2 from the *apo*-FabF2 crystal structure (PDB: 2GFW). ***h***, RMSFs – determined from simulations – of *apo*-FabF_2_* (apo structure derived from AcpP_2_=FabF_2_) onto loops 1 and 2 of the crosslinked AcpP_2_-FabF_2_ crystal structure. **i**, RMSFs determined from simulations of acyl-AcpP2•FabF2 mapped onto loops 1 and 2 of the crosslinked AcpP_2_-FabF_2_ crystal structure. In panels ***d***-***i***, larger (per residue) backbone RMSF values correspond to a thicker “sausage”; color range (blue to white to red) illustrates (per residue) side chain RMSFs with the blue-to-red color range indicating small-to-large RMSF values.

## Discussion

KSs catalyze multi-step reactions that require the association and dissociation of two distinct AcpP-tethered substrates, involving the intermediacy of an acyl-enzyme adduct. Enzymes that catalyze such complex chemical transformations often implement gating mechanisms to control solvent access, exert mechanistic selectivity, and/or control reaction order.^30^ Zhang *et al.* showed that Phe400 of *Streptococcus pneumoniae* FabF (SpFabF) acts as a KS gating residue.^17^ The F400A mutant of SpFabF has reduced catalytic activity relative to wildtype for its native reaction, but efficiently produces triacetic acid lactone (TAL). TAL is a common shunt product of condensing enzymes that results from the mispriming and concomitant extension of malonyl-ACP. These results suggest a compromised gating system that no longer controls reaction order. Additionally, ligand-bound and acyl-enzyme intermediate KS structures clearly show Phe400 in an alternate conformation, one more suitable for binding malonyl-ACP.^34,41^

We propose that elongating KSs use a double-drawbridge gating mechanism^30^ to direct substrate binding and control the timing and sequence of the transacylation and condensation half-reactions. In our model (Figure 6), ACP binds to *apo-*KS, upon which ACP’s helices 2 and 3 separate allowing the acyl cargo to chain-flip^25^ into the KS active site. KS loops 1 and 2 then move in a coordinated manner to assume a gate-open conformation forming a transient, hydrophobic delivery/extraction pocket that accommodates the substrate (Figure 6b). Loops 1 and 2 then return to their closed conformations, directing the substrate’s acyl chain into the fatty acyl binding pocket while simultaneously restoring the oxyanion hole that facilitates transacylation (Figure 6c). Upon transacylation, Phe400 rotates to form the malonyl-ACP binding pocket^22^ (Figure 6d). Malonyl-ACP then binds and undergoes a decarboxylative condensation reaction with the acyl-KS adduct to form β-ketoacyl-ACP. Loops 1 and 2 must then return to an open conformation to enable product dissociation.

**Figure 6.**
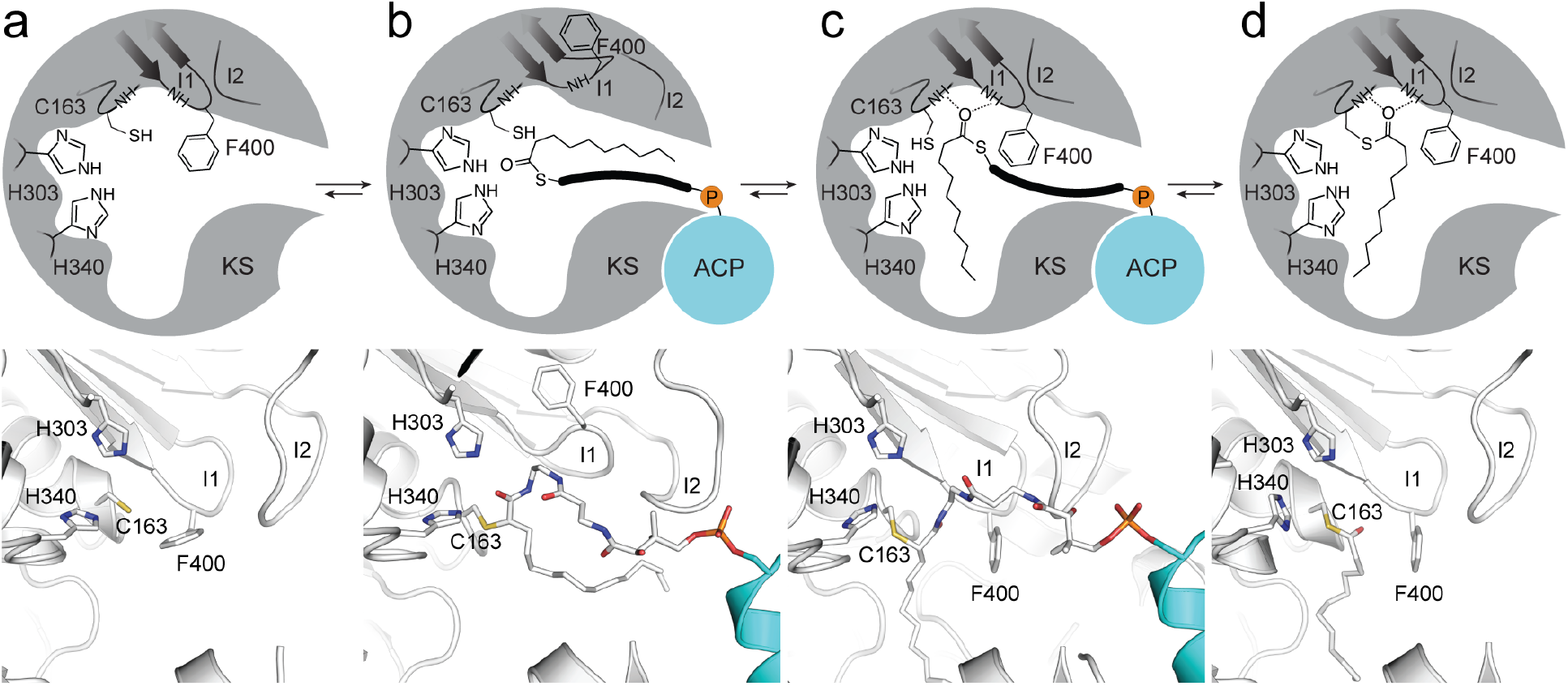
Proposed gating mechanism of elongating ketosynthases. Figure demonstrates the active site conformational changes that facilitate acyl-chain transfer in elongating KSs as demonstrated by trapped crystallographic states from this work and previous studies. ***a**,* The *apo-* form of the KS active site illustrated by a 2D schematic of the active site architecture (top) and the crystal structure of apo-FabF (PDB: 2GFW) (bottom). **b**, Active site reorganization to the “gate-open” conformation upon association of acyl-AcpP with KSs as seen in the C16AcpP-FabF structure reported herein with a 2D schematic of the active site (top) and pymol rendering of the active site (bottom). ***c**,* Active site of C12AcpP-FabB, illustrating the “gate-closed” transacylation competent form of the active sites of elongating KSs shown as 2D schematic (top) and pymol rendering (bottom). ***d**,* The acyl-enzyme adduct form of FabF (PDB: 2GFY) shown as a 2D schematic (top) and pymol rendering (bottom). All steps are represented with equilibrium arrows and we presume that the active site undergoes similar transformations to allow the abstraction of acyl group after condensation with malonyl-AcpP, whereby loop1 and loop2 will be required to return to their open states to facilitate removal and sequestration of the extended substrate from the KS active site to the hydrophobic core of ACP. Active site catalytic residues Cys163, His303, His340, and Phe400 are provided in FabF residue numbering, represented as sticks, and colored according to element (C-white, O-red, N-blue, S-yellow). Hydrogen-bonding interactions are represented as dotted lines.

The KASII class of condensing enzymes (e.g. FabF) are the evolutionary progenitors of KSs found in type I FASs and type I/II PKSs.^42,43^ Comparison of the sequences of FabF and the heterodimeric actinorhodin KS-chain length factor (KS-CLF), a representative KS from type II PKSs ^44^, shows that the GFGG gating motif (loop 1) is conserved in the KS but not in the catalytically inactive CLF (Figure S8c,d). Analysis of the KS domains in type I systems show that the loop 1 gating residues are present in the KS domain of human type I FAS as is the Asp265 of loop 2 (Figure S8c,d). Furthermore, the KS domains found in 6-deoxyerythronolide B synthase (DEBS), a representative of modular type I PKSs^45^, also retain loop 1 but have an altered sequence, GI/VSG instead of GFGG (Figure S8c,d). These differences may promote binding of branched (2*S*)-methylmalonyl-ACP extender units as the isoleucine or valine residues are less bulky than phenylalanine. Asp265 and Asn404, two highly conserved gatekeeping residues in FabF, are also present in DEBS; However, the remainder of loop 2 is poorly conserved, which may reflect different requirements for substrate processing. Despite these differences, a S315A mutation in loop 2 of DEBS KS1 increased the formation of propionate, which results from the decarboxylation of the methylmalonyl-ACP substrate.^46^ These findings suggest an impaired gating system, like that of the F400A mutant of SpFabF, that can no longer coordinate substrate transfer and C-C bond formation in the correct order. While gatekeeping elements appear to be a general feature of condensing enzymes based upon primary sequence analysis, a more precise stereochemical understanding of these elements will require further biochemical, structural, and computational interrogation.

Early work by Vagelos, Cronan, and Rock established differences in activity and substrate specificity for the *E. coli* elongating KSs, FabB and FabF.^36–38,47^ *In vivo*, FabB, in conjunction with the dehydratase FabA, produces unsaturated fatty acids from *de novo* FAB.^29,47–49^ Additionally, Cronan and coworkers demonstrated that FabF regulates membrane fluidity in response to changes in temperature through the conversion of *cis*-palmitoleoyl-AcpP (16:1-AcpP) into *cis*-vaccenoyl-AcpP (18:1-AcpP) at lower temperatures *in vivo* and *in vitro*, thereby increasing the amount of unsaturated fatty acids present in the membrane.^38,50^ Work to better delineate the role of FabB and FabF in *de novo* unsaturated FAB and thermal regulation of membrane fluidity is ongoing.

The importance of KS catalyzed carbon-carbon bond formation in natural product pathways cannot be overstated. These enzymes perform complex reactions and employ poorly understood biochemical regulatory mechanisms to ensure substrate specificity and product fidelity. Chemical, structural, and computational biology, together, uncovered the gating mechanisms used by KSs to recognize and process acyl-ACP substrates. These results provide broadly applicable insights into KS activity, function, and substrate selectivity as well as further our understanding of the *E. coli* AcpP interactome.

## Online Methods

### Experimental Methods

#### Ketosynthase recombinant proteins purification and thrombin cleavage

The N-terminal His_8_-tag FabF and FabB recombinant proteins were expressed in *E. coli* BL21, and grown in Terrific Broth. Cells were grown in the presence of 50 mg•L^−1^ kanamycin, induced with 0.5 mM IPTG at OD_600_ = 0.8, and incubated at 37 °C for 4 h. The cells were spun down by centrifugation at 7,000 rpm for 15 min and the collected pellets were resuspended in lysis buffer (50 mM Tris, 150 mM NaCl, 10% glycerol, pH 8.0) along with 0.5 mg•mL^−1^ lysozyme (Worthington Biochemical Corp). The pelleted cells were lysed by sonication (4 s pulses for 5 min), followed by another centrifugation at 18,000 rpm for 30 min to clear the lysate. The proteins were purified using Ni-NTA resin (Thermo) in the column and were washed sequentially with wash buffer (50 mM Tris, 150 mM NaCl, 10% glycerol, pH 8.0) followed by elution with 250 mM buffered imidazole. The His_8_-tags of purified proteins were cleaved with bovine thrombin (2 U per 1 mg protein) for 16 h at 10 °C while dialyzing against the dialysis buffer. Resulting solutions were re-purified using Ni-NTA resin (Thermo) to trap the un-cleaved proteins. The FabF and FabB were further purified by FPLC using the HiLoad Superdex 200 (GE) size exclusion column. The eluted proteins were collected and concentrated to 2-4 mg•mL^−1^ using Amicon Ultra Centrifuge Filters (Millipore) with 10 kDa molecular weight cut off.

#### Native AcpP recombinant protein purification

Native AcpP recombinant protein was expressed in *E. coli* BL21 (DE3) and grown in Terrific Broth. Cells were grown at 37 °C in the presence of 100 mg•L–1 ampicillin, induced with 0.5 mM IPTG at OD_600_ = 0.8, and incubated at 18 °C for 16 h. The cells were spun down by centrifugation at 2,000 rpm for 30 min, and the collected pellets were resuspended in lysis buffer (50 mM Tris, 5% glycerol, pH 7.4) along with 0.1 mg•mL^−1^ lysozyme (Worthington Biochemical Corp). The pelleted cells were lysed by sonication (4 sec pulses for 5 min). Due to the high stability of AcpP, irrelevant proteins were precipitated by dripping in same volume of isopropanol into the lysate at the speed of 0.1 mL•sec^−1^.^51^ The resulting mixture was spun down by centrifugation (10000 rpm for 1 h) and the supernatant was injected directly into FPLC. AcpP was purified by HiTrap Q HP anion exchange chromatography column in 50 mM Tris buffer with a gradient of NaCl from 0 M to 1 M and the native AcpP was eluted around 0.3 M NaCl (aq). The eluted protein was collected and concentrated using Amicon Ultra Centrifuge Filters (Millipore) with 3 kDa molecular weight cut off.

#### *Holo*-AcpP apofication and *apo*-AcpP purification

The purified AcpP was in a 7:3 mixture of *holo* and *apo-*form. The phosphopantetheine moiety on *holo*-AcpP was removed by acyl carrier protein hydrolase (AcpH) in lysis buffer for 16 h. Final reaction concentrations: 5 mg•mL^−1^ AcpP mixture, 0.01 mg•mL–1 AcpH, 50 mM Tris (pH 7.4), 10% glycerol, 10 mM MgCl_2_, 5 mM MnCl_2_,0.25 % DTT. The resulting pure *apo*-AcpP, was determined by urea PAGE, and further purified by HiLoad Superdex 75 size exclusion column (FPLC) to remove AcpH. The eluted protein was collected and concentrated using Amicon Ultra Centrifuge Filters (Millipore) with 3 kDa molecular weight cut off.

#### *Apo*-AcpP loading and crypto-AcpP purification

All crosslinkers were loaded onto *apo*-AcpP using the so-called “one-pot” chemoenzymatic method.^52^ This method utilizes three CoA biosynthetic enzymes (CoaA, CoaD, CoaE) to form CoA analogues and a PPTase (Sfp) to load them onto *apo*-AcpP, resulting in *crypto*-AcpP. Final reaction concentrations: 1 mg•mL^−1^ *apo*-AcpP, 0.04 mg•mL^−1^ Sfp, 0.01 mg•mL^−1^ CoaA, 0.01 mg•mL^−1^ CoaD, 0.01 mg•mL^−1^ CoaE, 50 mM potassium phosphate (pH 7.2), 12.5 mM MgCl_2_, 1 mM DTT, 0.2 mM crosslinker, 8 mM ATP. The stock solution of crosslinkers was prepared by dissolving them in DMSO to a final concentration of 100 mM for octanoate (C8) and dodecanoate (C12), and 50 mM for hexadecanoate (C16). Reactions were incubated under 37 °C for 16 h and then purified by HiLoad Superdex 75 (GE) size exclusion column. The eluted protein was collected and concentrated using Amicon Ultra Centrifuge Filters (Millipore) with 3 kDa molecular weight cut off up to a concentration about 1-3 mg•mL^−1^.

#### Crosslinking reaction and crosslinked complex purification

The crosslinking reactions were carried out by mixing *crypto*-AcpP with its partner protein (FabF or FabB) in 5:1 ratio at 37 °C for 16 h. Reactions were examined by 12 % SDS PAGE and purified by HiLoad Superdex 200 (GE) size exclusion column using minimal buffer (20 mM Tris, 50 mM NaCl, pH 7.4). The resulting protein complex was greater than 95 % in purity, determined by SDS PAGE, and was concentrated to 8-10 mg•mL^−1^ using Amicon Ultra Centrifuge Filters (Millipore) with 30 kDa molecular weight cut off. The concentrated crosslinked complex was immediately used for protein crystallization, otherwise it was flash-frozen and stored in –20 °C freezer for later experiments.

#### Crystallization, structure determination and refinement

The crystals of the crosslinked complexes were grown by v*apo*r diffusion in sitting drops at 6 °C. In detail, 1 μL of crosslinked complex (8-10 mg•mL^−1^) was mixed with 1 μL of corresponding buffer solution and the mixture was placed inverted over 500 μL of the well solution. The AcpP-FabF complexes crystallized in 26-30 % PEG 8K, 0.1 M sodium cacodylate pH 6.5, and 0.3 M sodium acetate, yielding numerous orange rods amongst a heavy precipitate. Seeding trials were performed to increase size and quality of the samples. The C12AcpP-FabB complexes produced crystals in two conditions: 20% PEG 8K, 0.2 M magnesium acetate, 0.1 M sodium cacodylate pH 6.5 as well as 30% PEG 8K, 0.2 M ammonium sulfate, 0.1 M sodium cacodylate pH 6.5. Both conditions yielded square plates and required one to two weeks were for complete growth of crystals. The C16AcpP-FabB complexes produced crystals in 18-24 % PEG 8K, 0.1 M sodium cacodylate pH 6.5, and 0.3 M sodium acetate.

All data were collected at the Advanced Light Source (ALS) synchrotron at Berkeley. Data were indexed using iMosflm then processed and scaled using the aimless program from the CCP4 software suite.^53,54^ Scaled reflection output data was used for model building in PHENIX.^55^ For AcpP-FabF complexes, phases were solved using molecular replacement by first locating the larger FabF monomer followed by a second search function to place the smaller AcpP molecule in the residual density. For (crosslinked) AcpP-FabB complexes, initial phases were solved using molecular replacement by first performing a search function using the FabB monomer and then manually placing the AcpP molecules in the residual electron density. The covalently attached 4’-phosphopantetheine parameter file was generated using Jligand (CCP4). Manually programmed parameter restraints were used to create the associated covalent bonds between 4’-phosphopantetheine to Ser36 and Cys163 during refinement.

## Computational Methods

### Simulation preparation

#### Preparation of apo-KS structures

Simulations of *apo*-FabB and *apo*-FabF were performed using two sets of initial coordinates: one derived from previously reported 1.55 Å *apo*-FabB (PDB: 2V69)^56^ and 2.40 Å *apo*-FabF (PDB: 2GFW)^57^ and a second set generated by modifying the crosslinked structures of *E. coli* AcpP-FabB and AcpP-FabF reported herein by deleting coordinates corresponding to atoms from AcpP.

### Preparation of acyl-AcpP•KS structures

The crosslinked structures of *E. coli* AcpP-FabB and AcpP-FabF reported herein were used to prepare the initial coordinates used to perform molecular dynamics. The asymmetric unit of the AcpP-FabF structure is – unlike that of crosslinked AcpP-FabB – is the 1:1 AcpP•FabF complex. In order to prepare coordinates for the (functional) biological unit, AcpP_2_•FabF_2_, the asymmetric unit of the AcpP_1_-FabF_1_ structure was rotated about the twofold symmetry axis of the AcpP=FabF biological unit. The 2:2 complexes of acyl-AcpPs shown in Scheme S1 and either FabB or FabF were constructed by manually modifying the chemical crosslinker present the experimental structures. Modifications of this moiety were performed using Avogadro 1.1.1,^58,59^ Gaussview 5.0,^60^ and Pymol v.1.8.6.^61^. Schrodinger was used to add poorly resolved C- and N-terminal residues omitted in the dimeric AcpP•FabB and AcpP•FabF complexes. The protonation state of all titratable residues in the *apo*-KSs and acyl-AcpP•KS didomains were assigned using the H++ webserver, with the exception of the active site Cys163 of FabB and FabF; this residue was simulated in its deprotonated (thiolate) form.^62-65^ Histidine protonation states were inspected manually.

### Parameter generation for acyl-AcpP

For each simulated AcpP•KS complex, the acyl substrate, phosphopantetheine cofactor, and the conserved serine residue of AcpP to which the cofactor is affixed were treated as a single nonstandard residue. AMBER (ff14SB)^66^ and GAFF^67^ type force field parameters were assigned to the atoms of these nonstandard residues using ANTECHAMBER.^68^ Partial charges for all atoms in all nonstandard residues were determined using the RESP methodology^69^ (HF/6-31G*). All quantum calculations were performed using Gaussian 09.^70^

### Preparation of simulation cells

Proteins were solvated in a water box with TIP3P^71^ water molecules. Using TLEAP^72^, the simulation cell was constructed such its edges were placed so that closest proteinogenic atom was 10 Å away. TLEAP was used to neutralize and ‘salt’ the water box to mimic physiological conditions (0.15 M). Counterions (Na^+^ and Cl^−^) were also added to the simulation cell randomly.

### Simulation details

Amber16^72^ was used to perform all molecular dynamics (MD)^73-78^ simulations. All simulations were performed using the ff14SB^76^ and GAFF^77^ force fields. A 2 fs time-step was utilized via the SHAKE algorithm, which constrains all nonpolar bonds involving hydrogen atoms.^79^ Long-range electrostatic interactions were treated using the Particle Mesh Ewald (PME) method with a 10 Å cutoff for all non-bonded interactions.^80^ Both solvated protein complexes were energy minimized in a two-step fashion. In a first step, solvent molecules and counterions were allowed to relax, while all protein atoms were restrained using a harmonic potential (k = 500 kcal mol^−1^ Å^−2^). This geometry optimization was followed by an unrestrained energy minimization of the entire system. The thermal energy available at a physiological temperature of 310 K was slowly added to each system over the course of a 3.5 ns NVT ensemble simulation. The solvated complexes were then subjected to unbiased isobaric-isothermal (NPT) simulations for 25 ns in order to equilibrate the heated structures. Three independent 500 ns production MD (NPT ensemble) of each system were performed. For both NVT and NPT simulations, the Langevin thermostat (λ = 5.0 ps^−1^) was used to maintain temperature control.^81^ Pressure regulation in NPT simulations (target pressure of 1 atm) was achieved by isotropic position scaling of the simulation cell volume using a Berendsen thermostat.

### Analysis and visualization of simulation data

Analysis was performed using CPPTRAJ, PYTRAJ, a Python front-end for the CPPTRAJ analysis code, and MDTRAJ.^82^ Trajectories were visualized using VMD 1.9.1^83^ and Pymol v1.8.6.4^61^ and Pymol v2.2.^84^ Coordinate data was written to disk every 10 ps.

## Data Availability Statement

All required crystallographic models and data were submitted to the Worldwide Protein Data Bank (wwPDB, www.wwpdb.org) under the accession codes 6OKC (C12AcpP-FabB), 6OKF(C16AcpP-FabB), 60LT (C12AcpP-FabF), 6OKG (C16AcpP-FabF).

## Code Availability Statement

All code used to process molecular dynamics simulation data is available upon request.

## Acknowledgements

This research was supported by National Science Foundation Grant EEC-0813570 (to J.P.N.), NIH RO1 GM095970 (to M.D.B.), NIH T32 G832626 for support of J.T.M., NIH T32 GM112584 to T.G.G., and NIH K12GM068524 for T.D.D. Portions of the work were also funded by the Arthur and Julie Woodrow Chair at the Salk Institute (to J.P.N.) and the Howard Hughes Medical Institute (to J.P.N.). The authors thank Dr. Gordon Louie for assistance in x-ray data collection, processing and refinement and Marianne Bowman for assistance with protein crystallization.

## Author contributions

J.L.L., J.A.M, M.D.B., and J.P.N. designed and supervised the project. T.D.D. synthesized probe molecules. J.T.M, W.E.K., T.G.B, A.C purified protein, optimized crosslinking, carried out crystallization, and obtained crystals. J.T.M. collected x-ray diffraction data and refined and analyzed crystallographic models. A.P. performed molecular dynamics simulations. A.P., J.T.M, and J.A.M analyzed simulation data. J.T.M, A.P., J.L.L., J.A.M, M.D.B., and J.P.N. wrote the manuscript. All authors edited the manuscript. Authors W.E.K., T.D.D., A.C, and T.G.B contributed equally to this work.

## Additional information

Chemical synthetic methods, chemical compound characterization, copies of NMR data on synthetic intermediates and source data for the X-ray crystal structures are available in the Supplementary Information.

## References

1. Hertweck, C. The biosynthetic logic of polyketide diversity. Angew. Chemie Int. Ed. 48, 4688–4716 (2009).

2. Staunton, J. & Weissman, K. J. Polyketide biosynthesis: a millennium review. Nat. Prod. Rep. 18, 380–416 (2001).

3. Beld, J., Lee, D. J. & Burkart, M. D. Fatty acid biosynthesis revisited: structure elucidation and metabolic engineering. Mol. Biosyst. 11, 38–59 (2015).

4. Ruppe, A. & Fox, J. M. Analysis of interdependent kinetic controls of fatty acid synthases. ACS Catal. 8, 11722–11734 (2018).

5. Cronan, J. E. & Thomas, J. Bacterial fatty acid synthesis and its relationships with polyketide synthetic pathways. Methods Enzymol. 459, 395–433 (2009).

6. Crosby, J. & Crump, M. P. The structural role of the carrier protein – active controller or passive carrier. Nat. Prod. Rep. 29, 1111–1137 (2012).

7. Chen, A., Re, R. N. & Burkart, M. D. Type II fatty acid and polyketide synthases: deciphering protein–protein and protein–substrate interactions. Nat. Prod. Rep. 35, 1029–1045 (2018).

8. Kapur, S. et al. Reprogramming a module of the 6-deoxyerythronolide B synthase for iterative chain elongation. Proc. Natl. Acad. Sci. 109, 4110 LP–4115 (2012).

9. Masoudi, A., Raetz, C. R. H., Zhou, P. & Pemble IV, C. W. Chasing acyl carrier protein through a catalytic cycle of lipid A production. Nature 505, 422 (2013).

10. Nguyen, C. et al. Trapping the dynamic acyl carrier protein in fatty acid biosynthesis. Nature 505, 427–431 (2014).

11. Lowry, B., Li, X., Robbins, T., Cane, D. E. & Khosla, C. A turnstile mechanism for the controlled growth of biosynthetic intermediates on assembly line polyketide synthases. ACS Cent. Sci. 2, 14–20 (2016).

12. Jiménez-Osés, G. et al. The role of distant mutations and allosteric regulation on LovD active site dynamics. Nat. Chem. Biol. 10, 431 (2014).

13. Blatti, J. L. et al. Manipulating fatty acid biosynthesis in microalgae for biofuel through protein-protein interactions. PLoS One 7, 1–12 (2012).

14. Chen, Y. et al. Structural classification and properties of ketoacyl synthases. Protein Sci. 20, 1659–1667 (2011).

15. Davies, C., Heath, R. J., White, S. W. & Rock, C. O. The 1.8 Å crystal structure and active-site architecture of β-ketoacyl-acyl carrier protein synthase III (FabH) from Escherichia coli. Structure 8, 185–195 (2000).

16. White, S. W., Zheng, J., Zhang, Y.-M. & Rock, C. O. The structural biology of type II fatty acid biosynthesis. Annu. Rev. Biochem. 74, 791–831 (2005).

17. Zhang, Y.-M., Hurlbert, J., White, S. W. & Rock, C. O. Roles of the active site water, histidine 303, and phenylalanine 396 in the catalytic mechanism of the elongation condensing enzyme of streptococcus pneumoniae. J. Biol. Chem. 281, 17390–17399 (2006).

18. von Wettstein-Knowles, P., Olsen, J. G., McGuire, K. A. & Henriksen, A. Fatty acid synthesis. FEBS J. 273, 695–710 (2006).

19. Zhang, Y.-M., Wu, B., Zheng, J. & Rock, C. O. Key residues responsible for acyl carrier protein and β-ketoacyl-acyl carrier protein reductase (FabG) interaction. J. Biol. Chem. 278, 52935–52943 (2003).

20. Zhang, Y.-M. et al. Identification and analysis of the acyl carrier protein (ACP) docking site on β-ketoacyl-ACP synthase III. J. Biol. Chem. 276, 8231–8238 (2001).

21. Tallorin, L. et al. Trapping of the enoyl-acyl carrier protein reductase-acyl carrier protein interaction. J. Am. Chem. Soc. 138, (2016).

22. Yu, X., Liu, T., Zhu, F. & Khosla, C. In vitro reconstitution and steady-state analysis of the fatty acid synthase from Escherichia coli. Proc. Natl. Acad. Sci. 108, 18643 LP–18648 (2011).

23. Dodge, G. J. et al. Structural and dynamical rationale for fatty acid unsaturation in *Escherichia coli* Proc. Natl. Acad. Sci. 201818686 (2019). doi:10.1073/pnas.1818686116

24. Butland, G. et al. Interaction network containing conserved and essential protein complexes in Escherichia coli. Nature 433, 531 (2005).

25. Cronan, J. E. The chain-flipping mechanism of ACP (acyl carrier protein)-dependent enzymes appears universal. Biochem. J. 460, 157 LP–163 (2014).

26. Sezonov, G., Joseleau-Petit, D. & D’Ari, R. Escherichia coli physiology in luria-bertani broth. J. Bacteriol. 189, 8746 LP–8749 (2007).

27. Jenner, M. et al. Acyl-chain elongation drives ketosynthase substrate selectivity in trans-acyltransferase polyketide synthases. Angew. Chemie Int. Ed. 54, 1817–1821 (2015).

28. Watanabe, K., Wang, C. C. C., Boddy, C. N., Cane, D. E. & Khosla, C. Understanding substrate specificity of polyketide synthase modules by generating hybrid multimodular synthases. J. Biol. Chem. 278, 42020–42026 (2003).

29. Magnuson, K., Jackowski, S., Rock, C. O. & Cronan, J. E. Regulation of fatty acid biosynthesis in Escherichia coli. Microbiol. Rev. 57, 522 LP–542 (1993).

30. Gora, A., Brezovsky, J. & Damborsky, J. Gates of enzymes. Chem. Rev. 113, 5871–5923 (2013).

31. Worthington, A. S., Porter, D. F. & Burkart, M. D. Mechanism-based crosslinking as a gauge for functional interaction of modular synthases. Org. Biomol. Chem. 8, 1769–1772 (2010).

32. Worthington, A. S. & Burkart, M. D. One-pot chemo-enzymatic synthesis of reporter-modified proteins. Org. Biomol. Chem. 4, 44–46 (2006).

33. Finzel, K., Lee, D. J. & Burkart, M. D. Using modern tools to probe the structure–function relationship of fatty acid synthases. ChemBioChem 16, 528–547 (2015).

34. Wang, J. et al. Platensimycin is a selective FabF inhibitor with potent antibiotic properties. Nature 441, 358–361 (2006).

35. Zondlo, N. J. Aromatic-Proline Interactions: Electronically Tunable CH/π Interactions. Acc. Chem. Res. 46, 1039–1049 (2013).

36. D’Agnolo, G., Rosenfeld, I. S. & Vagelos, P. R. Multiple forms of beta-ketoacyl-acyl carrier protein synthetase in Escherichia coli. J. Biol. Chem. 250, 5289–5294 (1975).

37. Garwin, J., Klages, A. & Cronan, J. Structural, enzymatic, and genetic studies of beta-ketoacyl-acyl carrier protein synthases I and II of Escherichia coli. J. Biol. Chem. 255, 11949–11956 (1980).

38. de Mendoza, D. & Cronan, J. E. Thermal regulation of membrane lipid fluidity in bacteria. Trends Biochem. Sci. 8, 49–52 (1983).

39. Edwards, P., Sabo Nelsen, J., Metz, J. G. & Dehesh, K. Cloning of the fabF gene in an expression vector and in vitro characterization of recombinant fabF and fabB encoded enzymes from Escherichia coli. FEBS Lett. 402, 62–66 (1997).

40. Henzler-Wildman, K. & Kern, D. Dynamic personalities of proteins. Nature 450, 964 (2007).

41. Trajtenberg, F. et al. Structural insights into bacterial resistance to cerulenin. FEBS J. 281, 2324–2338 (2014).

42. Jenke-Kodama, H., Sandmann, A., Müller, R. & Dittmann, E. Evolutionary implications of bacterial polyketide synthases. Mol. Biol. Evol. 22, 2027–2039 (2005).

43. Ridley, C. P., Lee, H. Y. & Khosla, C. Evolution of polyketide synthases in bacteria. Proc. Natl. Acad. Sci. U. S. A. 105, 4595–4600 (2008).

44. Keatinge-Clay, A. T., Maltby, D. A., Medzihradszky, K. F., Khosla, C. & Stroud, R. M. An antibiotic factory caught in action. Nat. Struct. & Mol. Biol. 11, 888 (2004).

45. Robbins, T., Liu, Y. C., Cane, D. E. & Khosla, C. Structure and mechanism of assembly line polyketide synthases. Curr. Opin. Struct. Biol. 41, 10–18 (2016).

46. Robbins, T., Kapilivsky, J., Cane, D. E. & Khosla, C. Roles of conserved active site residues in the ketosynthase domain of an assembly line polyketide synthase. Biochemistry 55, 4476–4484 (2016).

47. Heath, R. J. & Rock, C. O. Roles of the FabA and FabZ beta-hydroxyacyl-acyl carrier protein dehydratases in Escherichia coli fatty acid biosynthesis. J Biol Chem 271, (1996).

48. Feng, Y. & Cronan, J. E. Escherichia coli unsaturated fatty acid synthesis: Complex transcription of the FabA gene and in vivo identification of the essential reaction catalyzed by FabB. J. Biol. Chem. 284, 29526–29535 (2009).

49. Bloch, K. in (ed. Boyer, P. D. B. T.) 5, 441–464 (Academic Press, 1971).

50. Garwin, J. L. & Cronan, J. E. Thermal modulation of fatty acid synthesis in Escherichia coli does not involve de novo enzyme synthesis. J. Bacteriol. 141, 1457–1459 (1980).

