## Supplemental Information for "Gating Mechanism of β-Ketoacyl-ACP Synthases"

### Table of Contents

Figure S1. Chemistry and biology of ketosynthases. 3

Figure S2. Comparison of the protein-protein interactions between crosslinked AcpP=FabF and AcpP=FabB complexes. 5

Figure S3. Native contact analysis of MD simulations data of acyl-$AcpP_2$•$FabF_2$ (10:0-$AcpP_2$•$FabF_2$, 12:0-$AcpP_2$•$FabF_2$, 16:0-$AcpP_2$•$FabF_2$). 6

Figure S4. Native contact analysis of MD simulations data of acyl-$AcpP_2$•$FabB_2$ (10:0-$AcpP_2$•$FabB_2$, 12:0-$AcpP_2$•$FabB_2$, 16:0-$AcpP_2$•$FabB_2$). 7

Figure S5. Comparison of alkyl chain electron density and substrate•KS interactions in C16AcpP=FabF, C12AcpP=FabB, and C16AcpP=FabB crystal structures. 8

Figure S6. Analysis of contacts between active site residues and substrate of acyl-AcpP•FabF complexes sampled computationally. 10

Figure S7. Analysis of contacts between active site residues and substrate of acyl-AcpP•FabB complexes sampled computationally. 12

Figure S8. Analysis of loop 1 and 2 conservation by multi-sequence alignment (MSA) 13

Figure S9. Ramachandran plots of the GFGG motifs of FabB or FabF simulated in various states 14

Figure S10. Ramachandran plots of the key residues of loop 2 of FabB or FabF simulated in various states 16

Figure S11. Root mean square (RMS) fluctuations each residue of loops 1 and 2 of FabB and FabF monomers of the apo-AcpP2•KS2 sampled during MD simulations. 18

Figure S12. Root mean square (RMS) fluctuations each residue of loop 1 of FabB and FabF monomers of the acyl-AcpP2•KS2 sampled during MD simulations. 19

Figure S13. Root mean square (RMS) fluctuations each residue of loop 2 of FabB and FabF monomers of the acyl-AcpP2•KS2 sampled during MD simulations. 20

Electron density maps of reported structures. Stereo images of electron density maps for a. C16AcpP=FabF, (PDB: 6OKG, 2.4 Å). b. C12AcpP=FabB, (PDB: 6OKC ,1.55 Å) c. C16AcpP=FabB, (PDB: 6OKF, 2.5 Å.) 22

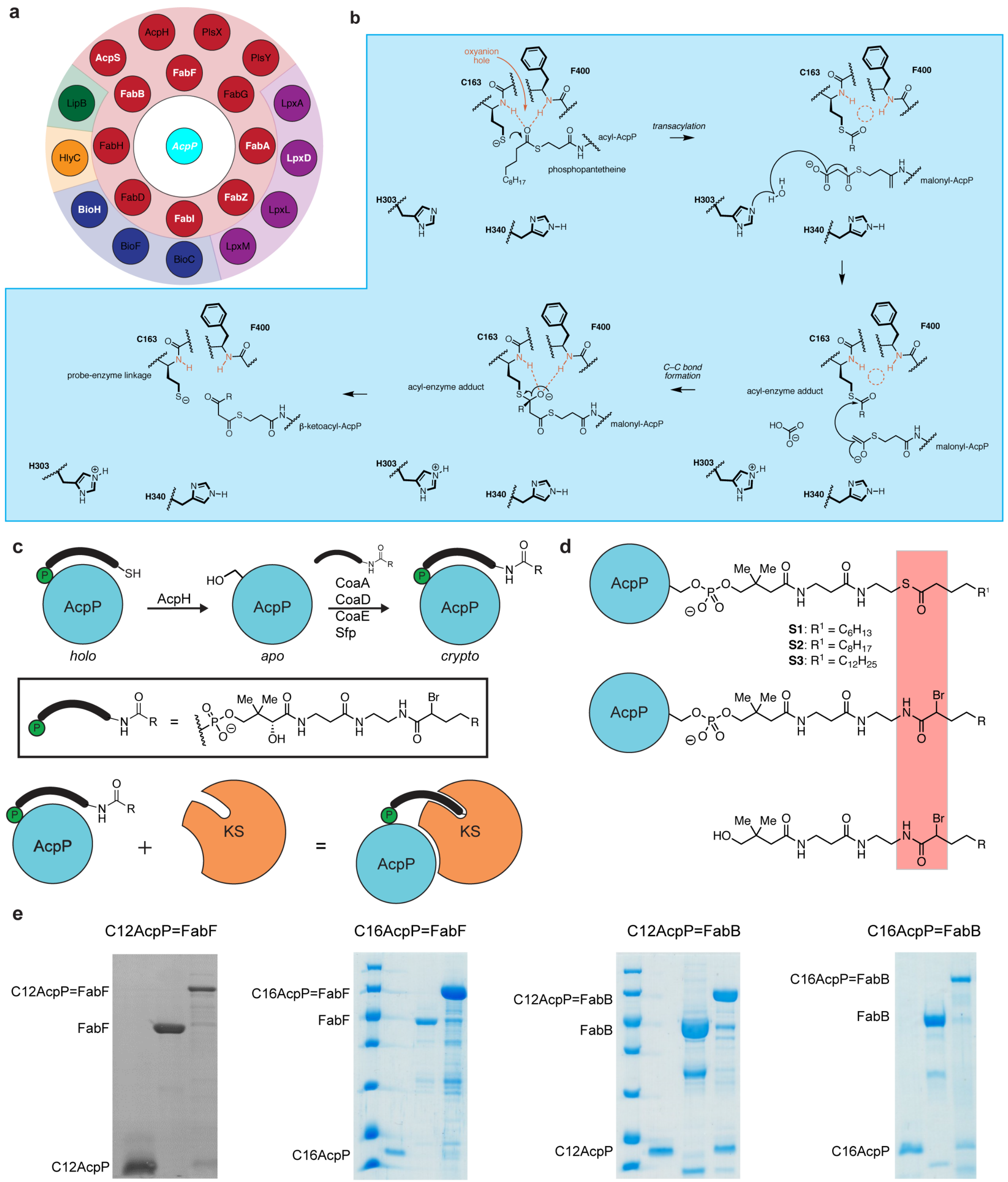


**Figure S1.** Chemistry and biology of ketosynthases. **a**. ACP "Interactome" in *E. coli*. Proteins are colored by biosynthetic pathway (green: fatty acid biosynthesis, purple: lipid A biosynthesis, Blue: biotin biosynthesis, yellow: hemolysin activation, red: lipoyl activation of lipoate-dependent enzymes. **b.** Proposed reaction mechanism of elongating ketoacyl-AcpP-synthase FabB (KASI) and FabF (KASII). Top row of mechanism shows the transacylation half-reaction, while the bottom portion illustrates the condensation half-reaction. **c.** Crosslinking workflow scheme demonstrating apofication and loading of ACP with crosslinking probes to produce *crypto*-ACP, which is then mixed with partner protein to produce a crosslinked complex. **d.** Comparison of acyl-AcpP to crypto-AcpP loaded with substrate or crosslinking substrate mimetic, respectively. **S1**, **S2**, and **S3** indicate the acyl-AcpPs subjected to simulations as constituents of acyl-AcpP•KS complexes reported herein **e.** Crosslinking gels of C12AcpP=FabF, C16AcpP=FabF, C12AcpP=FabB, and C16AcpP=FabB.

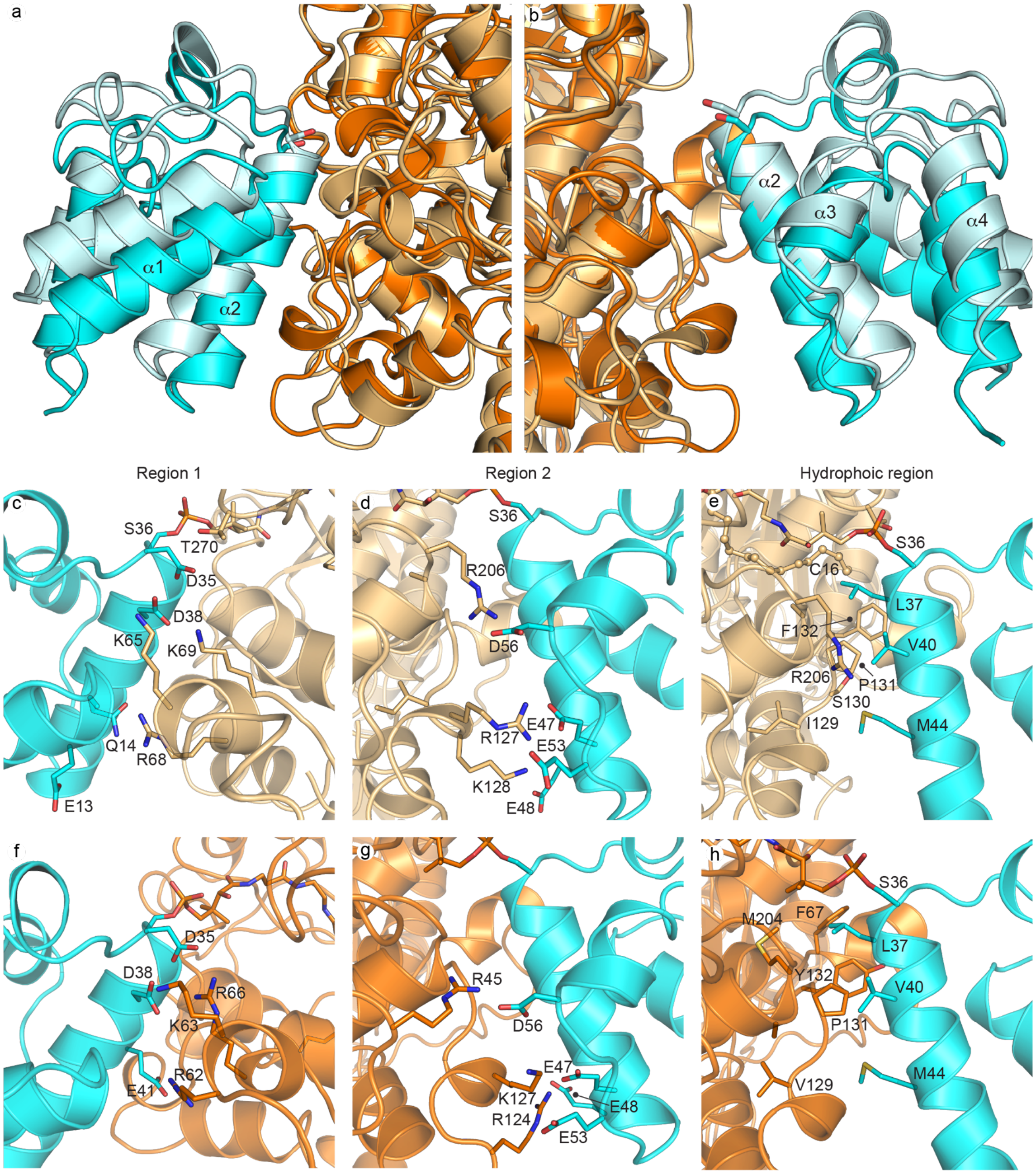


**Figure S2.** Comparison of the protein-protein interactions between crosslinked AcpP=FabF and AcpP=FabB complexes. AcpP is shown as cyan in all panels. FabF is shown in light orange and FabB is shown in bright orange. **a. b.** Superposition of C12AcpP=FabB (AcpP shown as light cyan) with C16AcpP=FabB (AcpP shown as cyan) from two 180° related views demonstrating differences in the angle and conformation of AcpP engagement with each KS interface. **c.** C16AcpP=FabF region 1 interface interactions **d.** C16AcpP=FabF region 2 interface interactions. **e.** C16AcpP=FabF hydrophobic interface interactions. **f.** C12AcpP=FabB region 1 interface interactions **g.** C12AcpP=FabB region 2 interface interactions. **h.** C12AcpP=FabB hydrophobic interface interactions

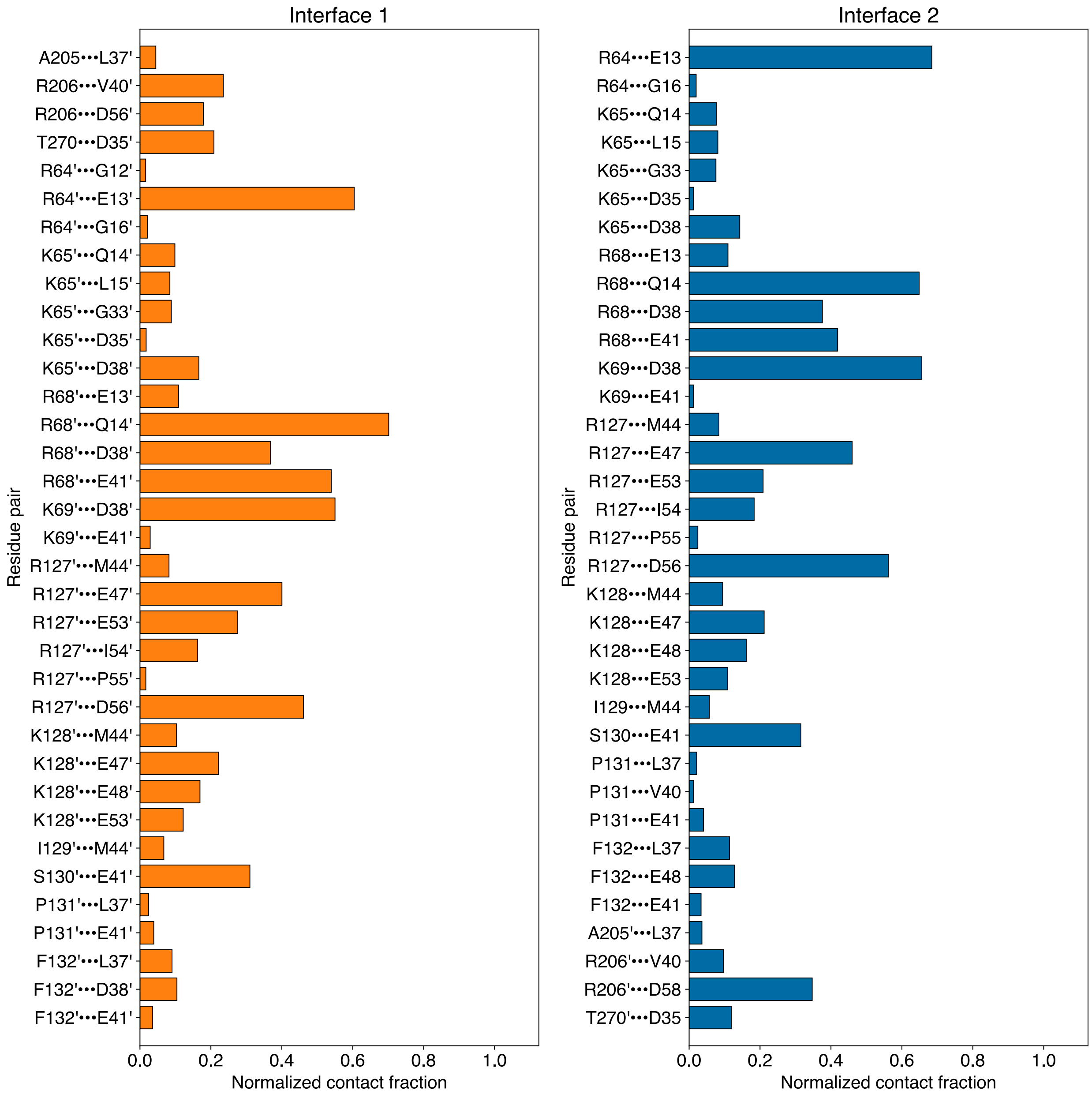


**Figure S3.** Native contact analysis of MD simulations data of acyl-AcpP$_2$•FabF$_2$ (10:0-AcpP$_2$•FabF$_2$, 12:0-AcpP$_2$•FabF$_2$, 16:0-AcpP$_2$•FabF$_2$). Residues on the dimeric FabF$_2$ subunit within 7.5 Å and those of either AcpP1 (Interface 1) and AcpP2 (Interface 2) within 7.5 Å dimer subunit were subjected to this analysis. The bar graphs illustrate the frequency with interfacial residues contact with one another at either interface of these acyl-AcpP$_2$•FabF$_2$. The contact fraction is defined as the total fraction of simulation data in which a residue pair is engaged in an intermolecular contact. A distance criterion of 3.5 Å or less between a pair of heavy atoms of two residue belonging two distinct protein monomers defines such a contact. This value before normalization assume values greater than 1.0, if a residue pair possesses multiple atoms that satisfy the distance criteria used to define a contact. Only pairwise contacts with contact fractions greater than or equal to 0.05 are included in the plots above. Every 10$^{th}$ frame of simulation data starting with the first was used to perform this analysis. Note that “primed” residues (e.g., K127’, D35’) refer to residues belonging to FabF2 or AcpP2.

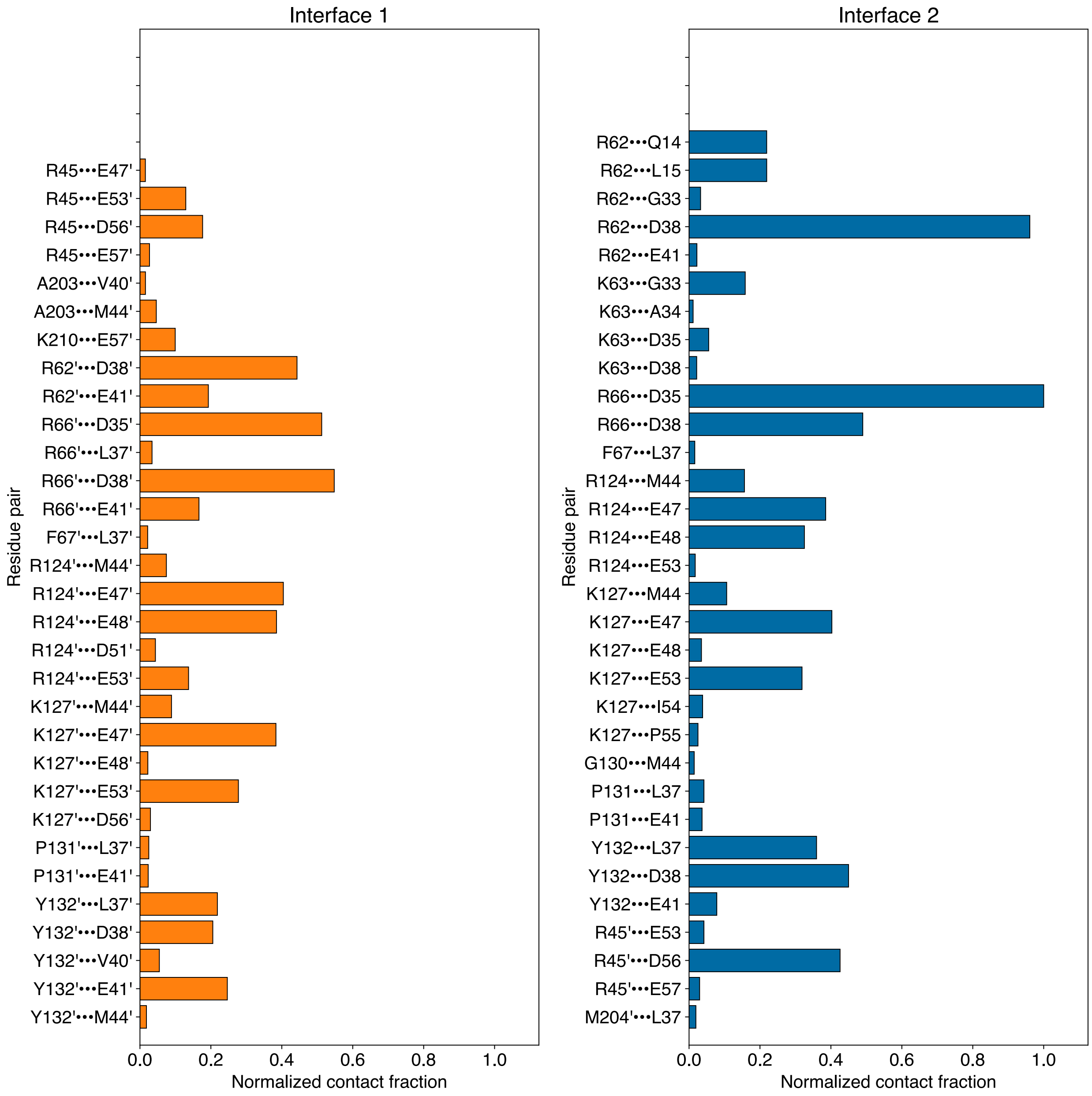


**Figure S4.** Native contact analysis of MD simulations data of acyl-AcpP$_2$•FabB$_2$ (10:0-AcpP$_2$•FabB$_2$, 12:0-AcpP$_2$•FabB$_2$, 16:0-AcpP$_2$•FabB$_2$). Residues on the dimeric FabB$_2$ subunit within 7.5 Å and those of either AcpP1 (Interface 1) and AcpP2 (Interface 2) within 7.5 Å dimer subunit were subjected to this analysis. The bar graphs illustrate the frequency with interfacial residues contact with one another at either interface of these acyl-AcpP$_2$•FabB$_2$. The contact fraction is defined as the total fraction of simulation data in which a residue pair is engaged in an intermolecular contact. A distance criterion of 3.5 Å or less between a pair of heavy atoms of two residue belonging two distinct protein monomers defines such a contact. This value before normalization assume values greater than 1.0, if a residue pair possesses multiple atoms that satisfy the distance criteria used to define a contact. Only pairwise contacts with contact fractions greater than or equal to 0.05 are included in the plots above. Every $10^{th}$ frame of simulation data starting with the first was used to perform this analysis. Note that “primed” residues (e.g., K127’, D35’) refer to residues belonging to FabB2 or AcpP2.

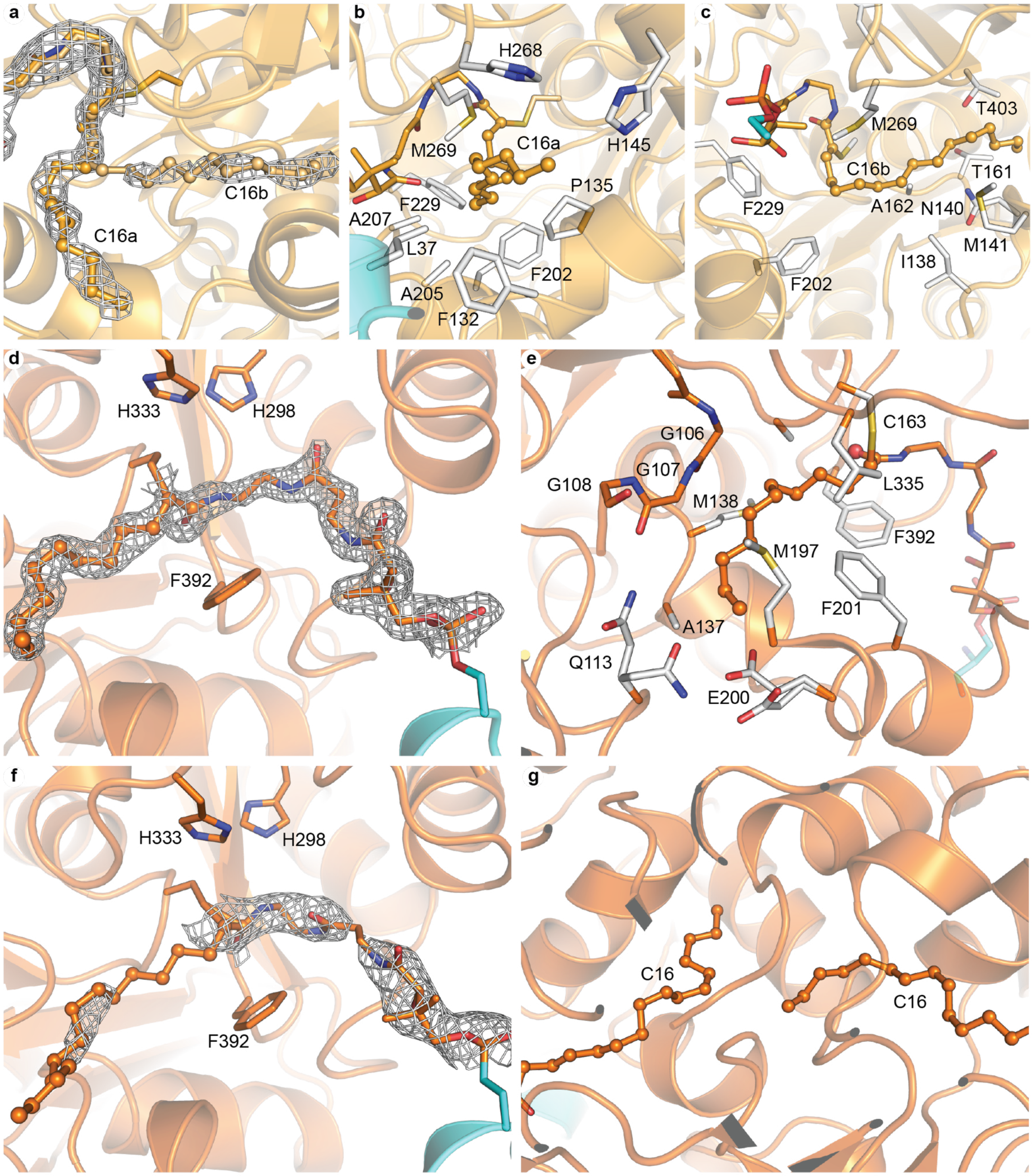


**Figure S5.** Comparison of alkyl chain electron density and substrate•KS interactions in C16AcpP=FabF, C12AcpP=FabB, and C16AcpP=FabB crystal structures. AcpP is shown as cyan in all panels. FabF is shown in light orange and FabB is shown in bright orange. **a.** Electron density of C16 alkyl chain in conformations a (C16a) and b (C16b) from the C16AcpP=FabF structure contoured with 2FO-FC maps at 1.0 σ **b.** All FabF residues within 4 Å of alkyl chain conformation a (C16a). **c.** All residues within 4 Å of alkyl chain conformation b (C16b). **d.** Electron density of C12 alkyl chain from C12AcpP=Fabb structure contoured at 1.0 σ. **e.** All FabB residues within 4 Å of the C12 alkyl chain substrate analog. **f.** Electron density of C16PPant probe from the C16AcpP=FabB structure contoured at 1.0 σ. **h.** Attempts to model a C16 probe reveal a constricted active site that places alkyl chains bound within the active site of both KS monomers in close proximity to one another and in unfavorable conformations.

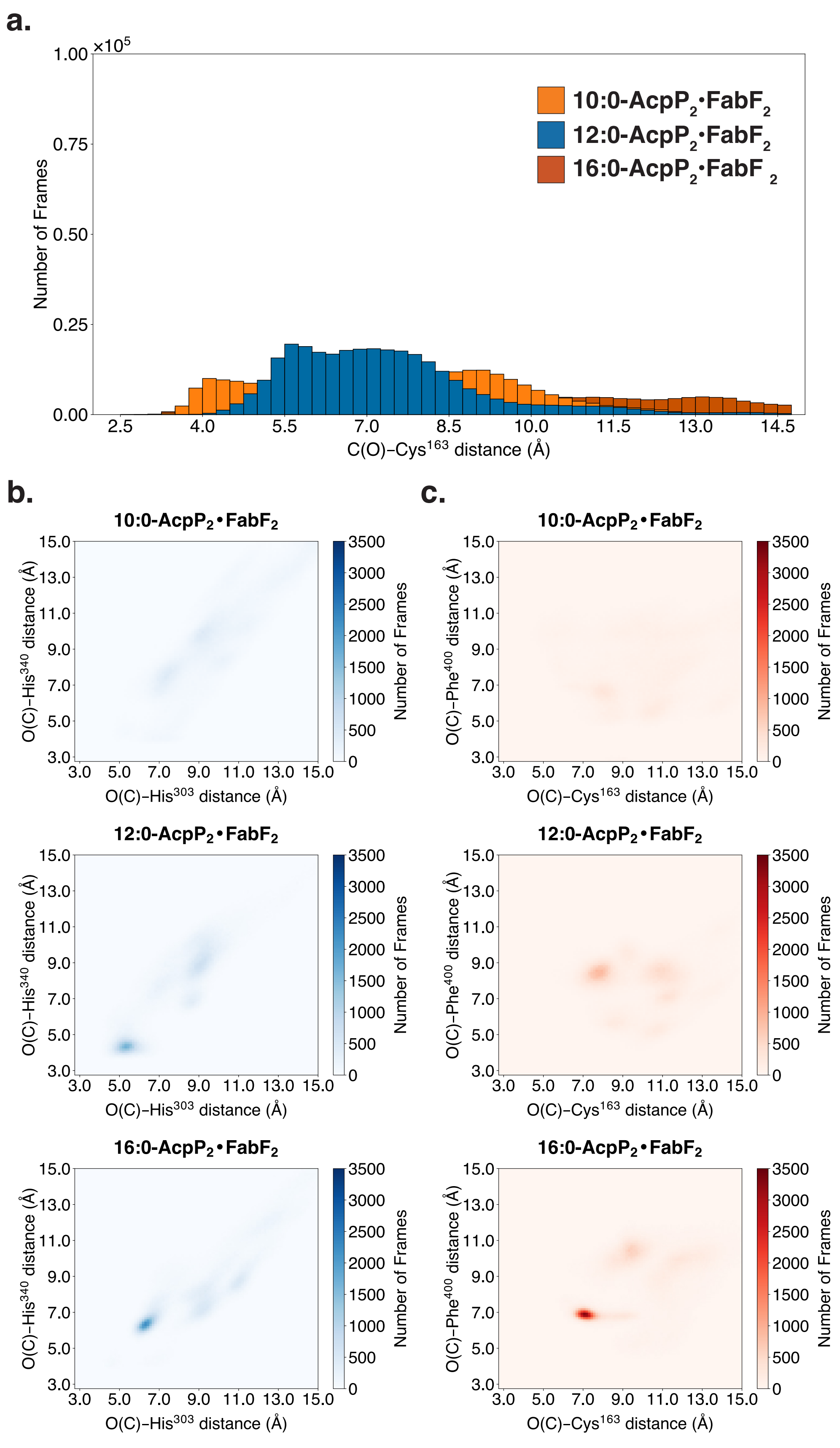

a.
10:0-AcpP₂•FabF₂
12:0-AcpP₂•FabF₂
16:0-AcpP₂•FabF₂
Number of Frames
C(O)–Cys163 distance (Å)
b.
c.
10:0-AcpP₂•FabF₂
12:0-AcpP₂•FabF₂
16:0-AcpP₂•FabF₂
O(C)–His340 distance (Å)
O(C)–His303 distance (Å)
O(C)–Phe400 distance (Å)
O(C)–Cys163 distance (Å)
Number of Frames

**Figure S6.** Analysis of contacts between active site residues and substrate of acyl-AcpP•FabF complexes sampled computationally. a. Distribution of distances sampled between carbonyl carbon of the acyl chain of 10:0-AcpP•FabF (orange), 12:0-AcpP-FabF (blue), or 16:0-AcpP•FabF (red) and sulfur atom of catalytic cysteine (Cys163). 1D histogram was generated using a bin width of 0.25 Å b. Analysis of hydrogen bonding interactions involving the substrate and His303 and His340 (2D histogram, blue). Simulation data is binned along two coordinates The first of which is the distance between the carbonyl oxygen of the substrate and the imidazole moiety of His303 (center of geometry) of FabF, while the second measures the distance between the imidazole moiety of His340 (center of geometry) and its carbonyl oxygen. c. Analysis of backbone (oxyanion hole) hydrogen-bonding interactions (2D histogram, red). Simulation data is binned along two coordinates. The first of which is the distance between the carbonyl oxygen of the substrate and the backbone amide nitrogen of Cys163 of FabF, while the second measures the distance between substrate's carbonyl oxygen and the backbone amide nitrogen of Phe400. Simulation data is binned along two coordinates. Data was sorted into 0.05 Å × 0.05 Å bins. Color bars indicate the absolute population of each histogram bin.

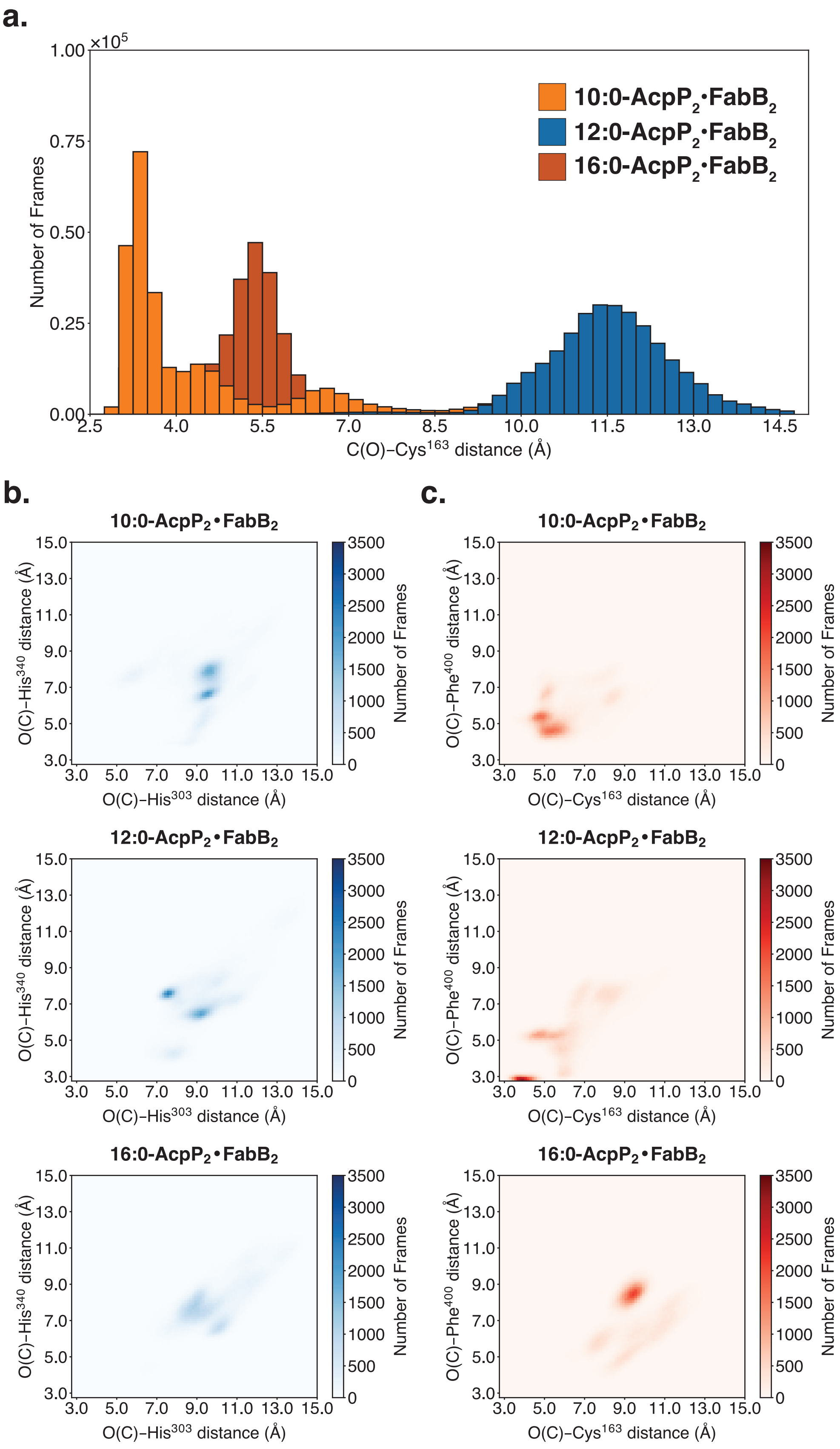

a.
×10^5
Number of Frames
C(O)–Cys163 distance (Å)
10:0-AcpP2•FabB2
12:0-AcpP2•FabB2
16:0-AcpP2•FabB2
b.
c.
10:0-AcpP2•FabB2
12:0-AcpP2•FabB2
16:0-AcpP2•FabB2
O(C)–His340 distance (Å)
O(C)–His303 distance (Å)
O(C)–Phe400 distance (Å)
O(C)–Cys163 distance (Å)
Number of Frames

**Figure S7.** Analysis of contacts between active site residues and substrate of acyl-AcpP•FabB complexes sampled computationally. a. Distribution of distances sampled between carbonyl carbon of the acyl chain of 10:0-AcpP•FabB (orange), 12:0-AcpP-FabB (blue), or 16:0-AcpP•FabB (red) and sulfur atom of catalytic cysteine (Cys163). 1D histogram was generated using a bin width of 0.25 Å b. Analysis of hydrogen bonding interactions involving the substrate and His303 and His340 (2D histogram, blue). Simulation data is binned along two coordinates The first of which is the distance between the carbonyl oxygen of the substrate and the imidazole moiety of His303 (center of geometry) of FabB, while the second measures the distance between the imidazole moiety of His340 (center of geometry) and its carbonyl oxygen. c. Analysis of backbone (oxyanion hole) hydrogen-bonding interactions (2D histogram, red) Simulation data is binned along two coordinates. The first of which is the distance between the carbonyl oxygen of the substrate and the backbone amide nitrogen of Cys163 of FabB, while the second measures the distance between substrate's carbonyl oxygen and the backbone amide nitrogen of Phe400. Simulation data is binned along two coordinates. Data was sorted into 0.05 Å × 0.05 Å bins. Color bars indicate the absolute population of each histogram bin.

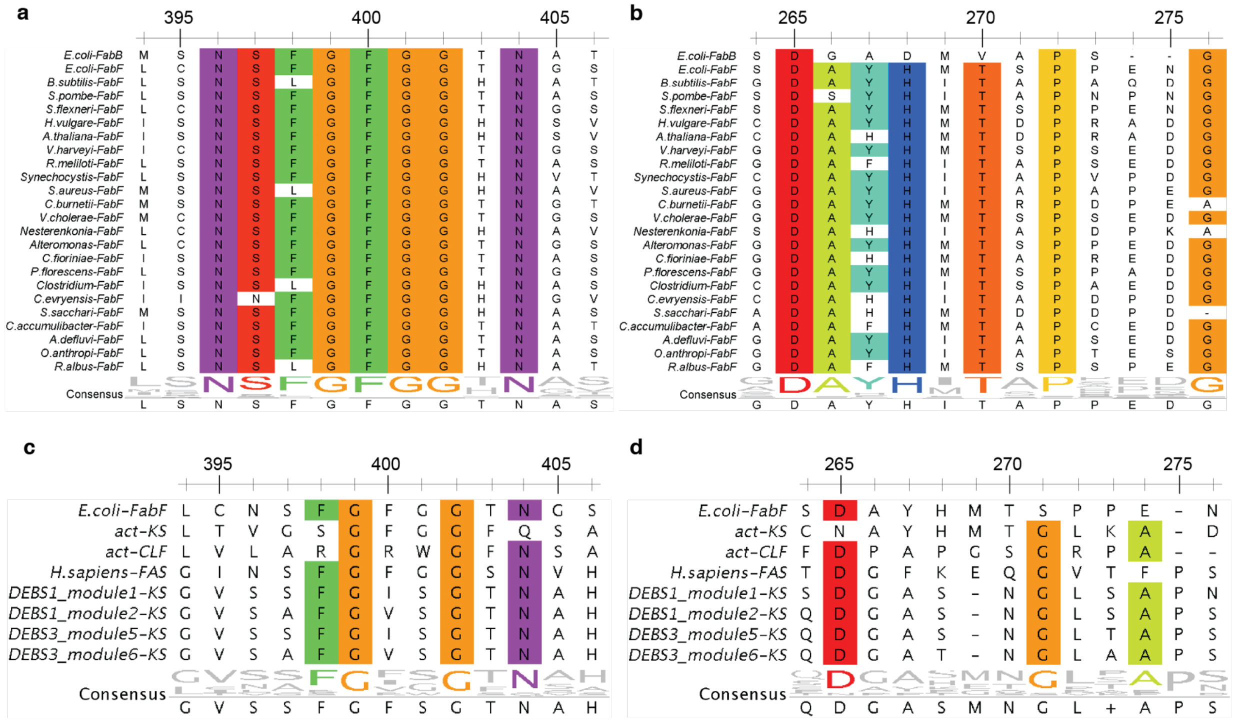


**Figure S8.** Analysis of loop 1 and 2 conservation by multi-sequence alignment (MSA) **a.** MSA alignment of FabB and 100 FabF orthologues comparing the Loop 1 GFGG β-turn region and surrounding conserved residues (FabF numbering). **b**. MSA Alignment of FabB and 100 FabF orthologues comparing the loop 2 sequence and surrounding conserved residues (FabF numbering). **c.** MSA comparison of FabF loop 1 with related ketosynthases from the actinorhodin biosynthetic (act-KS, act-CLF), human fatty acid biosynthesis (*H. sapiens*-FAS), and deoxyerythronolide B biosynthetic pathway (DEBS). **d.** MSA comparison of FabF loop 2 with related ketosynthases from the actinorhodin biosynthetic pathway, ketosynthase-chain length factor (KS-CLF) (act-KS, act-CLF), human fatty acid biosynthesis (*H. sapiens*-FAS), and deoxyerythronolide B biosynthetic pathway (DEBS). Sequences were aligned using the muscle algorithm in Mega7 and viewed using Jalview. Positions with greater than 70% identity were colored by the Taylor color scheme. The consensus sequence for the region is shown below the alignment.

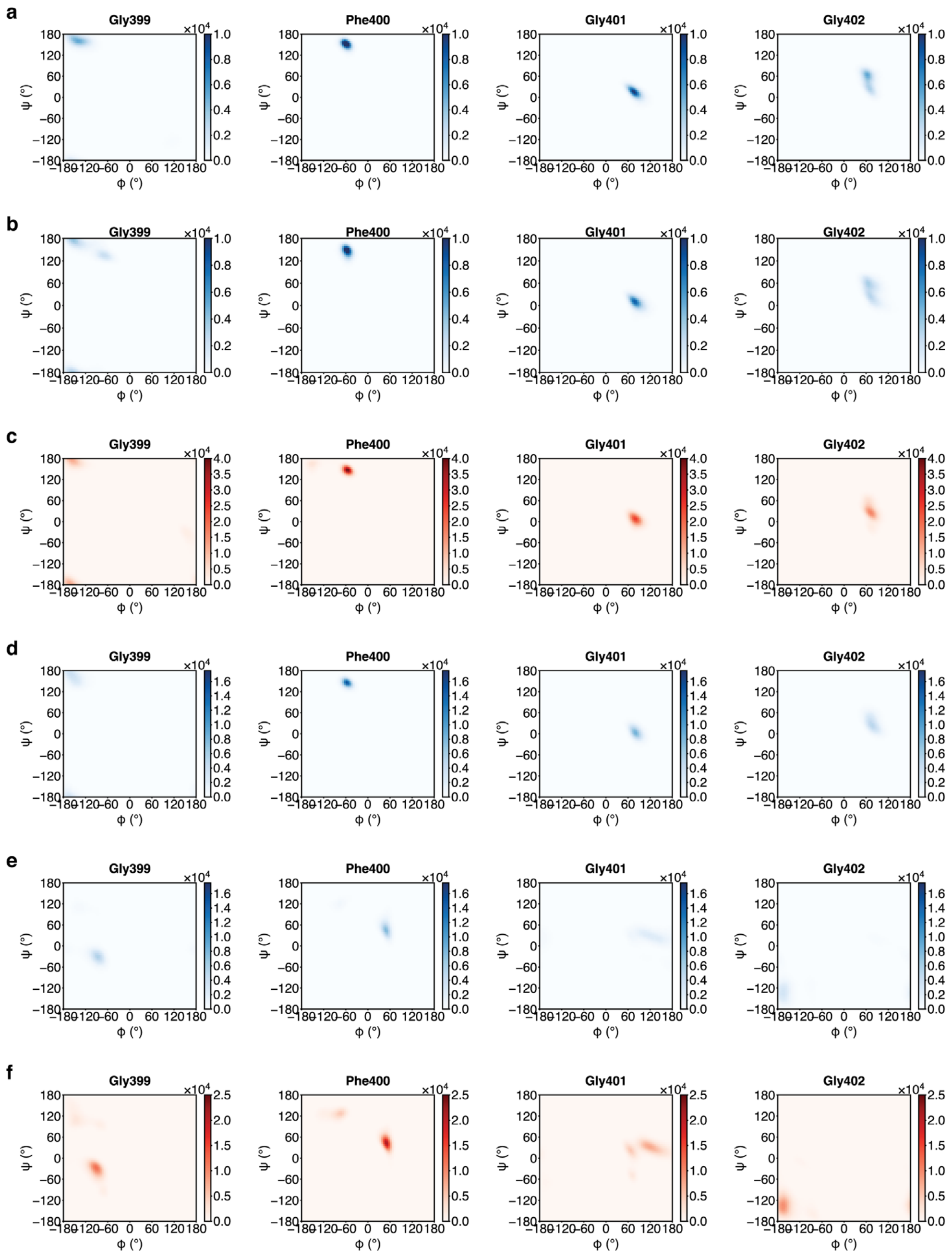


**Figure S9.** Ramachandran plots of the GFGG motifs of FabB or FabF simulated in various states. **a.** Analysis of the GFGG backbone dihedrals of *apo*-FabB performed using 1.5 µs of molecular dynamics (MD) simulation data of *apo*-FabB. **b.** Analysis of GFGG backbone dihedrals on apo-FabB* (apo-FabB constructed using the coordinates from the crosslinked AcpP=FabB structure reported herein) using 1.5 µs of molecular dynamics (MD) simulation data of *apo*-FabB*. **c.** Analysis of the GFGG backbone dihedral of acyl-AcpP•FabB using 4.5 µs of molecular dynamics (MD) simulation data of acyl-AcpP•FabB. **d.** Analysis of the GFGG backbone dihedrals of *apo*-FabF performed using 1.5 µs of molecular dynamics (MD) simulation data of *apo*-FabF. **e.** Analysis of GFGG backbone dihedrals on apo-FabF* (apo-FabF constructed using the coordinates from the crosslinked AcpP=FabF structure reported herein) using 1.5 µs of molecular dynamics (MD) simulation data of *apo*-FabF*. **f.** Analysis of the GFGG backbone dihedral of acyl-AcpP•FabF using 4.5 µs of molecular dynamics (MD) simulation data of acyl-AcpP•FabF. Data is binned based backbone ψ and ϕ dihedral angles residues assume during MD simulations using bin widths of 5°

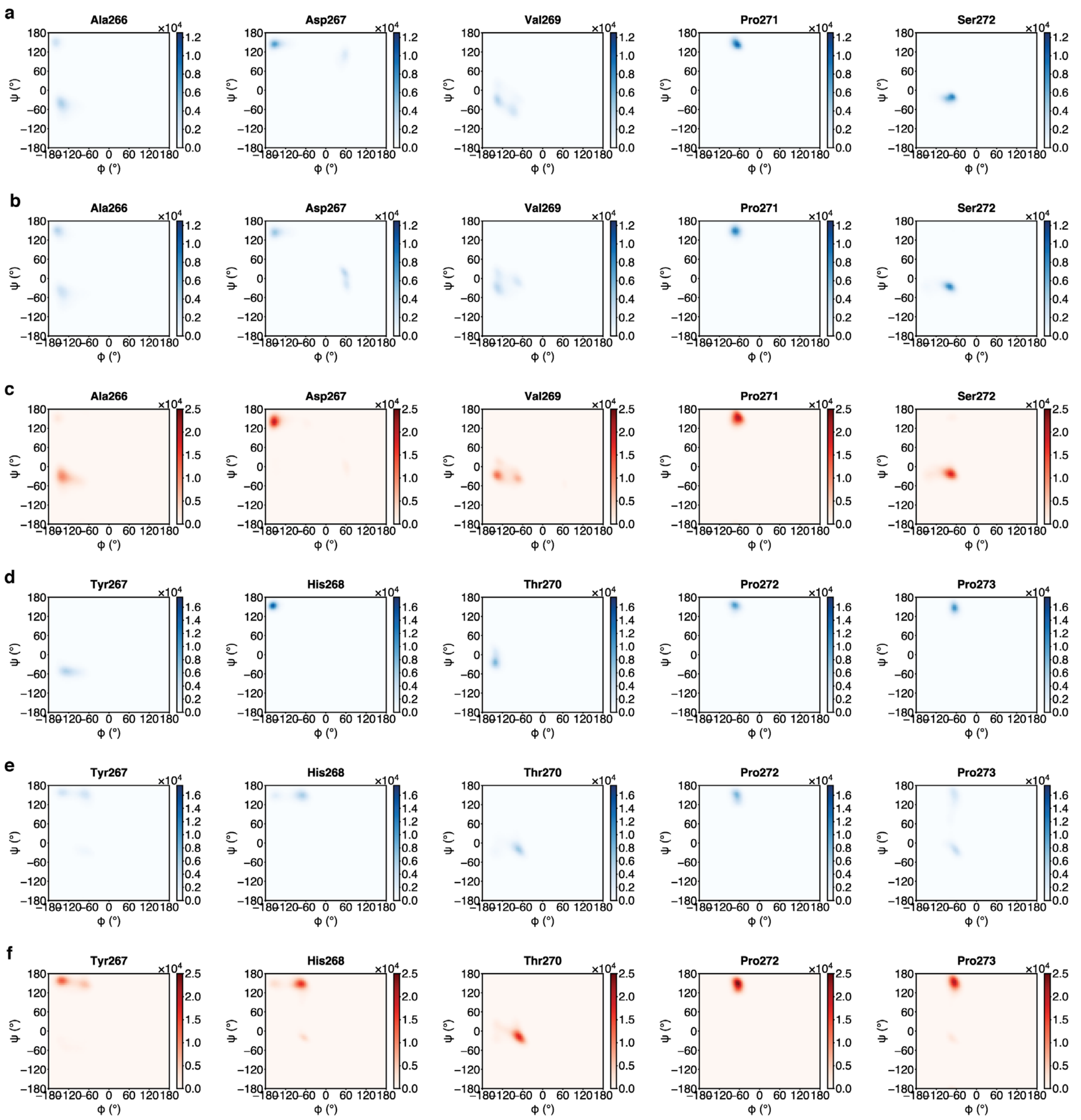


**Figure S10.** Ramachandran plots of the key residues of loop 2 of FabB or FabF simulated in various states. **a.** Analysis of the loop 2 backbone dihedrals of *apo*-FabB performed using 1.5 µs of molecular dynamics (MD) simulation data of *apo*-FabB. **b.** Analysis of loop 2 backbone dihedrals on apo-FabB* (apo-FabB constructed using the coordinates from the crosslinked AcpP=FabB structure reported herein) using 1.5 µs of molecular dynamics (MD) simulation data of *apo*-FabB*. **c.** Analysis of the loop 2 backbone dihedrals of acyl-AcpP•FabB using 4.5 µs of molecular dynamics (MD) simulation data of acyl-AcpP•FabB. **d.** Analysis of the loop 2 backbone dihedrals of *apo*-FabF performed using 1.5 µs of molecular dynamics (MD) simulation data of *apo*-FabB. **e.** Analysis of loop 2 backbone dihedrals on apo-FabF* (apo-FabF constructed using the coordinates from the crosslinked AcpP=FabF structure reported herein) using 1.5 µs of molecular dynamics (MD) simulation data of *apo*-FabB*. **f.** Analysis of the loop 2 backbone dihedral of acyl-AcpP•FabF using 4.5 µs of molecular dynamics (MD) simulation data of acyl-AcpP•FabF. Data is binned based backbone ψ and ϕ dihedral angles residues assume during MD simulations using bin widths of 5°.

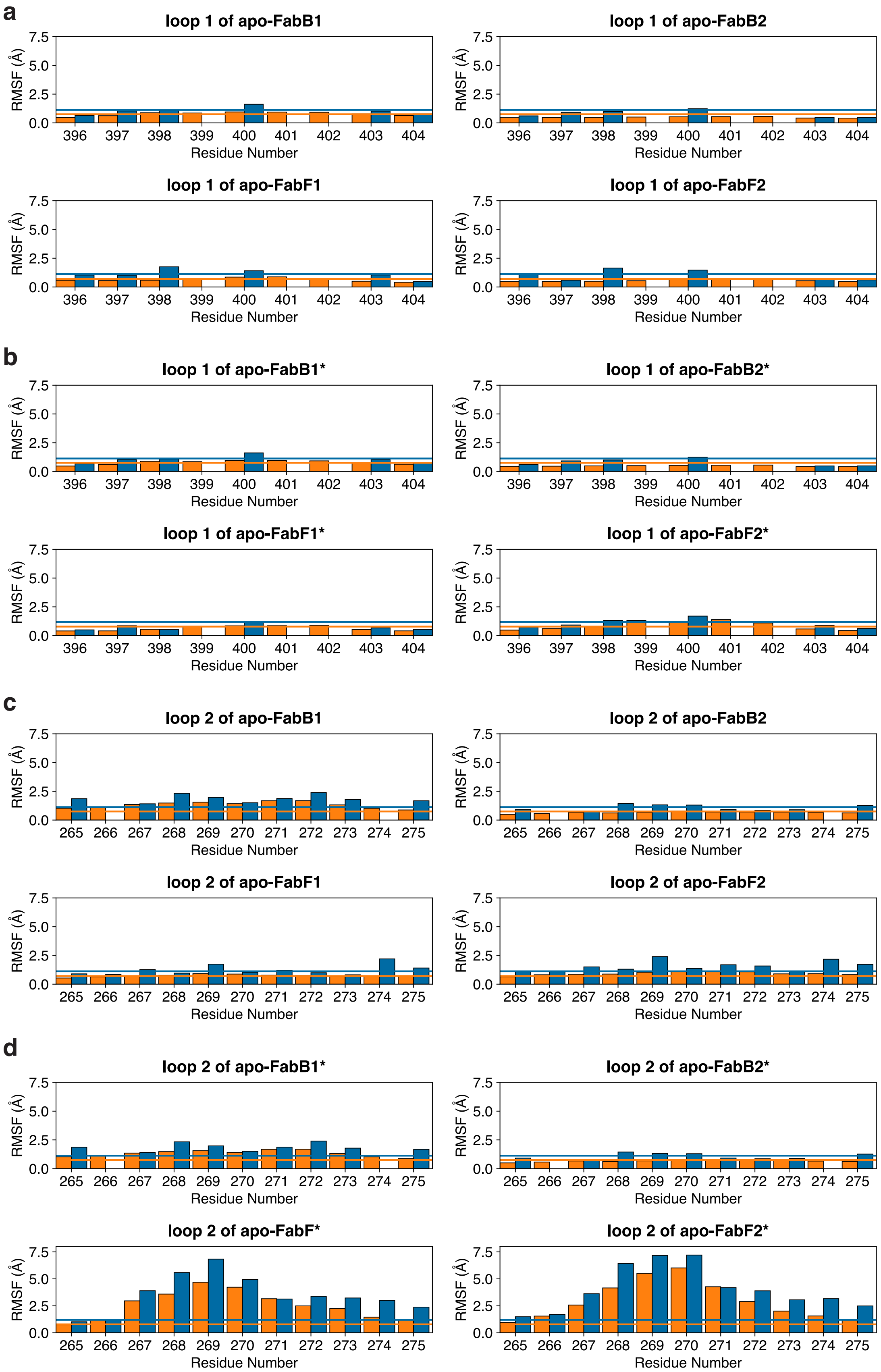
a
loop 1 of apo-FabB1
loop 1 of apo-FabB2
loop 1 of apo-FabF1
loop 1 of apo-FabF2
b
loop 1 of apo-FabB1*
loop 1 of apo-FabB2*
loop 1 of apo-FabF1*
loop 1 of apo-FabF2*
c
loop 2 of apo-FabB1
loop 2 of apo-FabB2
loop 2 of apo-FabF1
loop 2 of apo-FabF2
d
loop 2 of apo-FabB1*
loop 2 of apo-FabB2*
loop 2 of apo-FabF*
loop 2 of apo-FabF2*
RMSF (Å)
Residue Number
0.0
2.5
5.0
7.5
396 397 398 399 400 401 402 403 404
265 266 267 268 269 270 271 272 273 274 275

**Figure S11.** Root mean square (RMS) fluctuations each residue of loops 1 and 2 of FabB and FabF monomers of the *apo*-AcpP•KS sampled during MD simulations. **a.** Per residue RMSF of loop 1 of either monomer of *apo*-KS dimer. **b.** Per residue RMSF of loop 1 of either monomer of *apo*-KS dimer derived from the crosslinked AcpP=KS (*apo*-KS*) structure. **c.** Per residue RMSF of loop 2 of either monomer of *apo*-KS dimer. **d.** Per residue RMSF of loop 2 of either monomer of apo-*KS* dimer derived from the crosslinked AcpP=KS (*apo*-KS*) structure. The bar graphs show the per residue backbone RMS fluctuations (blue, measured using backbone heavy atoms) and the side chain RMS fluctuation of sidechains (orange, measured using each residue's sidechain heavy atoms). Residues 396-404 and residues 265-275 (FabF residue numbering) define loops 1 and 2, respectively.

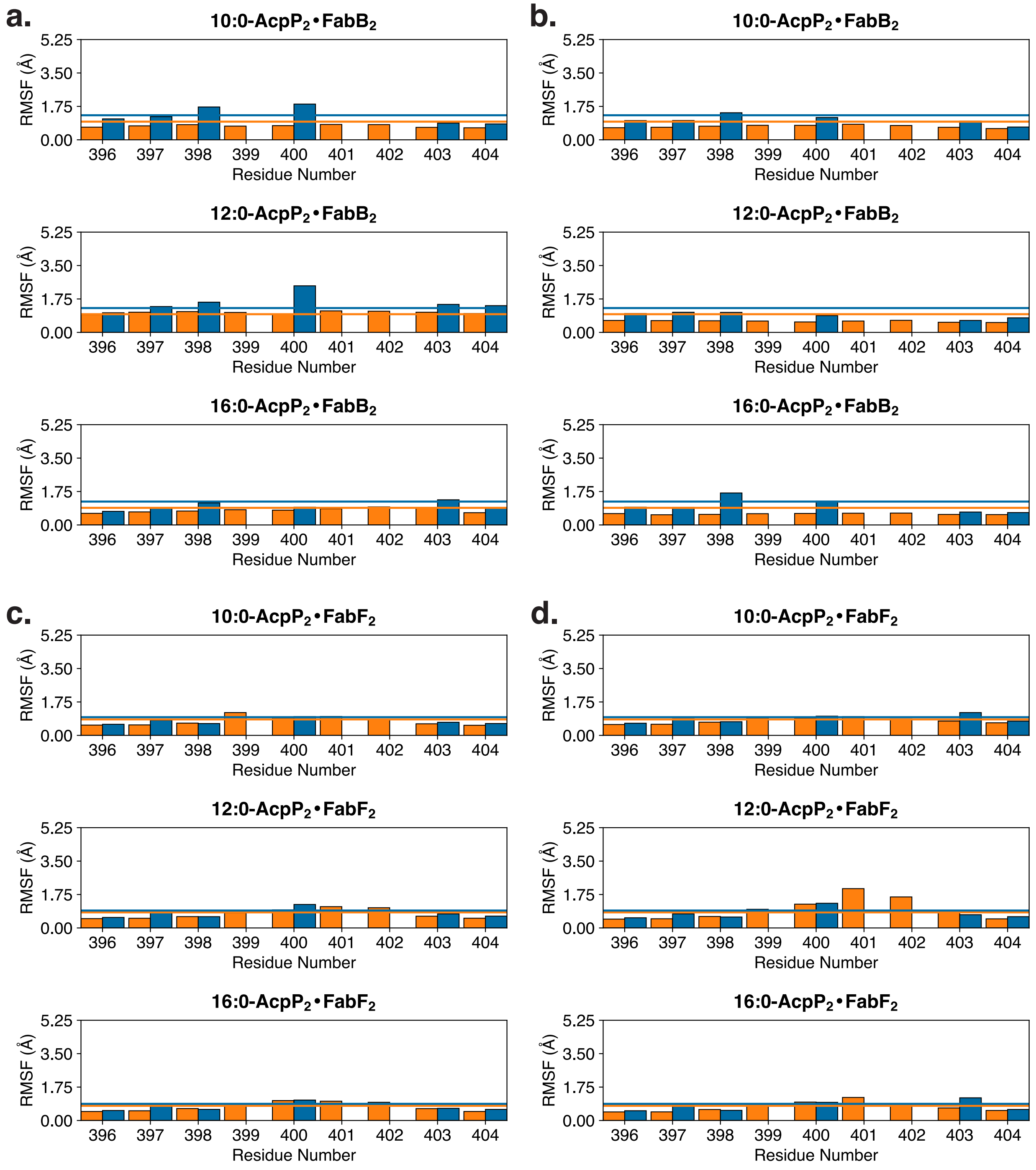


**Figure S12.** Root mean square (RMS) fluctuations each residue of loop 1 of FabB and FabF monomers of the acyl-AcpP•KS sampled during MD simulations. **a.** Per residue RMSF of loop 1 of the FabB1 monomer of acyl-AcpP•FabB. **b.** Per residue RMSF of loop 1 of the FabB2 monomer of acyl-AcpP•FabB. **c.** Per residue RMSF of loop 1 of the FabF1 monomer of acyl-AcpP•FabF. **d.** Per residue RMSF of loop 1 of the FabF2 monomer of acyl-AcpP•FabF. The bar graphs show the per residue backbone RMS fluctuations (blue, measured using backbone heavy atoms) and the side chain RMS fluctuation of sidechains (orange, measured using each residue's sidechain heavy atoms). Residues 396-404 (FabF residue numbering) define loop 1.

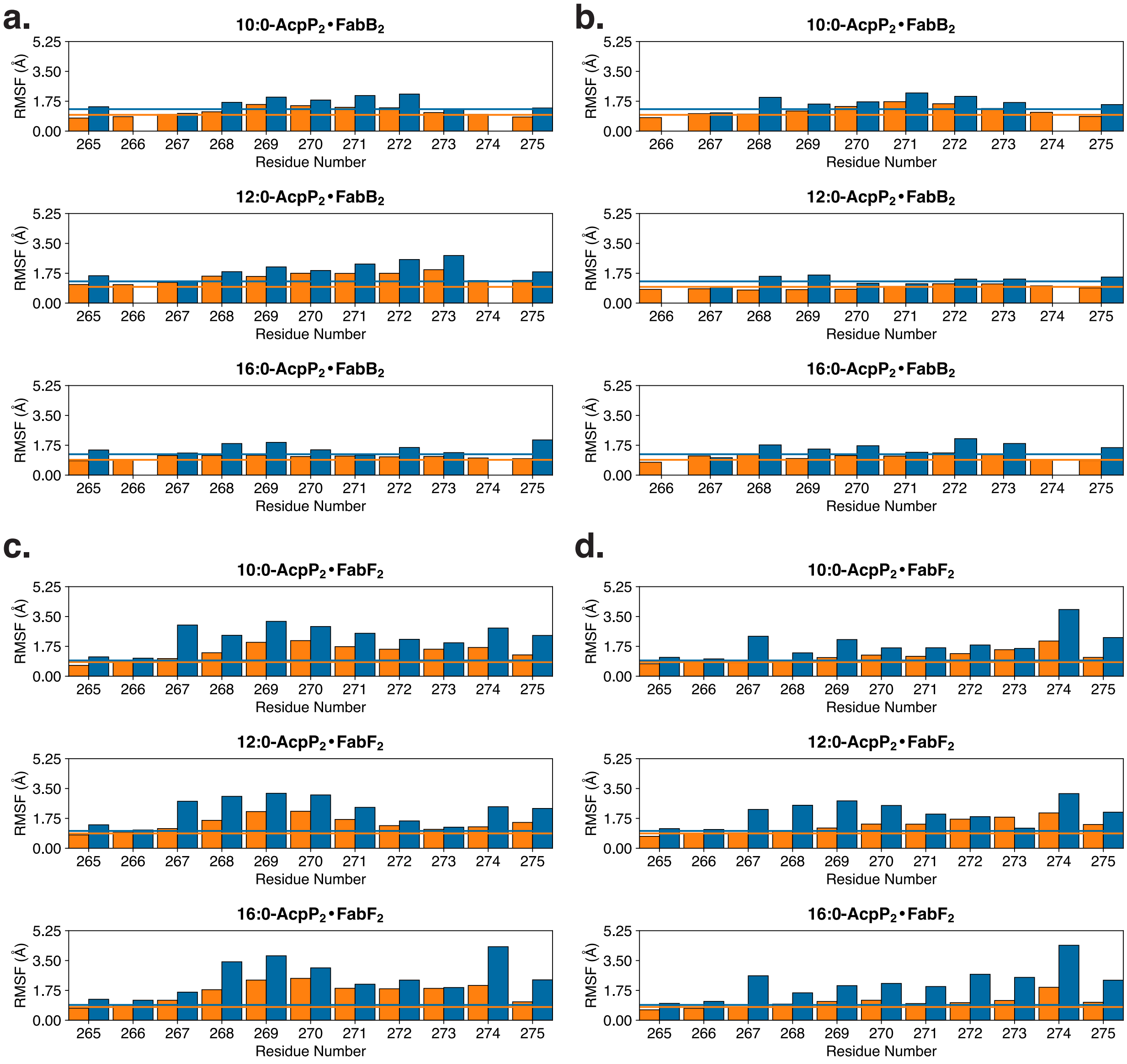


**Figure S13.** Root mean square (RMS) fluctuations each residue of loop 2 of FabB and FabF monomers of the acyl-AcpP2•KS2 sampled during MD simulations. **a.** Per residue RMSF of loop 2 of the FabB1 monomer of acyl-AcpP•FabB. **b.** Per residue RMSF of loop 2 of the FabB2 monomer of acyl-AcpP•FabB. **c.** Per residue RMSF of loop 2 of the FabF1 monomer of acyl-AcpP•FabF. **d.** Per residue RMSF of loop 2 of the FabF2 monomer of acyl-AcpP•FabF. The bar graphs show the per residue backbone RMS fluctuations (blue, measured using backbone heavy atoms) and the side chain RMS fluctuation of sidechains (orange, measured using each residue's sidechain heavy atoms). Residues 265-275 (FabF residue numbering) define loop 2.

**Table 1.** Refinement statistics from structures solved in this work.

| Table 1. Data Collection and Refinement Statistics | | | | |
|---|---|---|---|---|
| | **C16AcpP-FabB** | **C12AcpP-FabB** | **C16AcpP-FabF** | **C12AcpP-FabF** |
| **Wavelength** | | | | |
| **Resolution range** | 60.22 - 2.5 (2.589 - 2.5) | 87.7 - 1.549 (1.604 - 1.549) | 69.04 - 2.3 (2.382 - 2.3) | 69.14 - 2.35 (2.434 - 2.35) |
| **Space group** | P 1 21 1 | P 21 21 21 | P 43 21 2 | P 43 21 2 |
| **Unit cell** | 58.939 100.781 79.259 90 108.64 90 | 59.0704 112.17 140.68 90 90 90 | 86.8896 86.8896 113.729 90 90 90 | 86.4497 86.4497 115.16 90 90 90 |
| **Total reflections** | 186617 (19226) | 1593391 (155305) | 381910 (38754) | 288893 (29114) |
| **Unique reflections** | 30112 (2997) | 135556 (13249) | 19983 (1952) | 18796 (1830) |
| **Multiplicity** | 6.2 (6.4) | 11.8 (11.7) | 19.1 (19.9) | 15.4 (15.9) |
| **Completeness (%)** | 98.74 (98.19) | 99.43 (98.28) | 99.80 (99.95) | 99.74 (99.67) |
| **Mean I/sigma(I)** | 7.59 (2.34) | 14.25 (3.65) | 9.28 (1.70) | 9.91 (1.08) |
| **Wilson B-factor** | 45.37 | 13.45 | 27.25 | 40.96 |
| **R-merge** | 0.1373 (0.8075) | 0.1099 (0.8567) | 0.3746 (2.379) | 0.3274 (3.604) |
| **R-meas** | 0.15 (0.8784) | 0.1149 (0.8954) | 0.3848 (2.44) | 0.3385 (3.722) |
| **R-pim** | 0.05957 (0.3421) | 0.03317 (0.2574) | 0.0868 (0.5384) | 0.0852 (0.9233) |
| **CC1/2** | 0.993 (0.908) | 0.998 (0.906) | 0.977 (0.62) | 0.996 (0.45) |
| **CC*** | 0.998 (0.976) | 1 (0.975) | 0.994 (0.875) | 0.999 (0.788) |
| **Reflections used in refinement** | 30078 (2991) | 135501 (13236) | 19953 (1951) | 18770 (1829) |
| **Reflections used for R-free** | 1642 (159) | 6845 (675) | 997 (98) | 1094 (107) |
| **R-work** | 0.1785 (0.2673) | 0.1477 (0.1934) | 0.1946 (0.2811) | 0.2006 (0.3147) |
| **R-free** | 0.2175 (0.2890) | 0.1630 (0.2044) | 0.2279 (0.3078) | 0.2389 (0.2938) |
| **CC(work)** | 0.967 (0.904) | 0.972 (0.934) | 0.522 (0.780) | 0.961 (0.728) |
| **CC(free)** | 0.938 (0.864) | 0.953 (0.917) | 0.938 (0.733) | 0.944 (0.740) |
| **RMS(bonds)** | 0.004 | 0.008 | 0.004 | 0.004 |
| **RMS(angles)** | 0.99 | 1.3 | 0.97 | 0.96 |
| **Ramachandran favored (%)** | 94.86 | 96.65 | 96.9 | 95.67 |
| **Ramachandran allowed (%)** | 4.4 | 3.04 | 3.1 | 3.92 |
| **Ramachandran outliers (%)** | 0.73 | 0.31 | 0 | 0.41 |
| **Rotamer outliers (%)** | 7.09 | 1.8 | 1.05 | 5.94 |
| **Clashscore** | 5.48 | 2.84 | 2.86 | 6.69 |
| **Average B-factor** | 65.05 | 22.28 | 31.9 | 59.21 |
| **Number of TLS groups** | 19 | 34 | 11 | 12 |

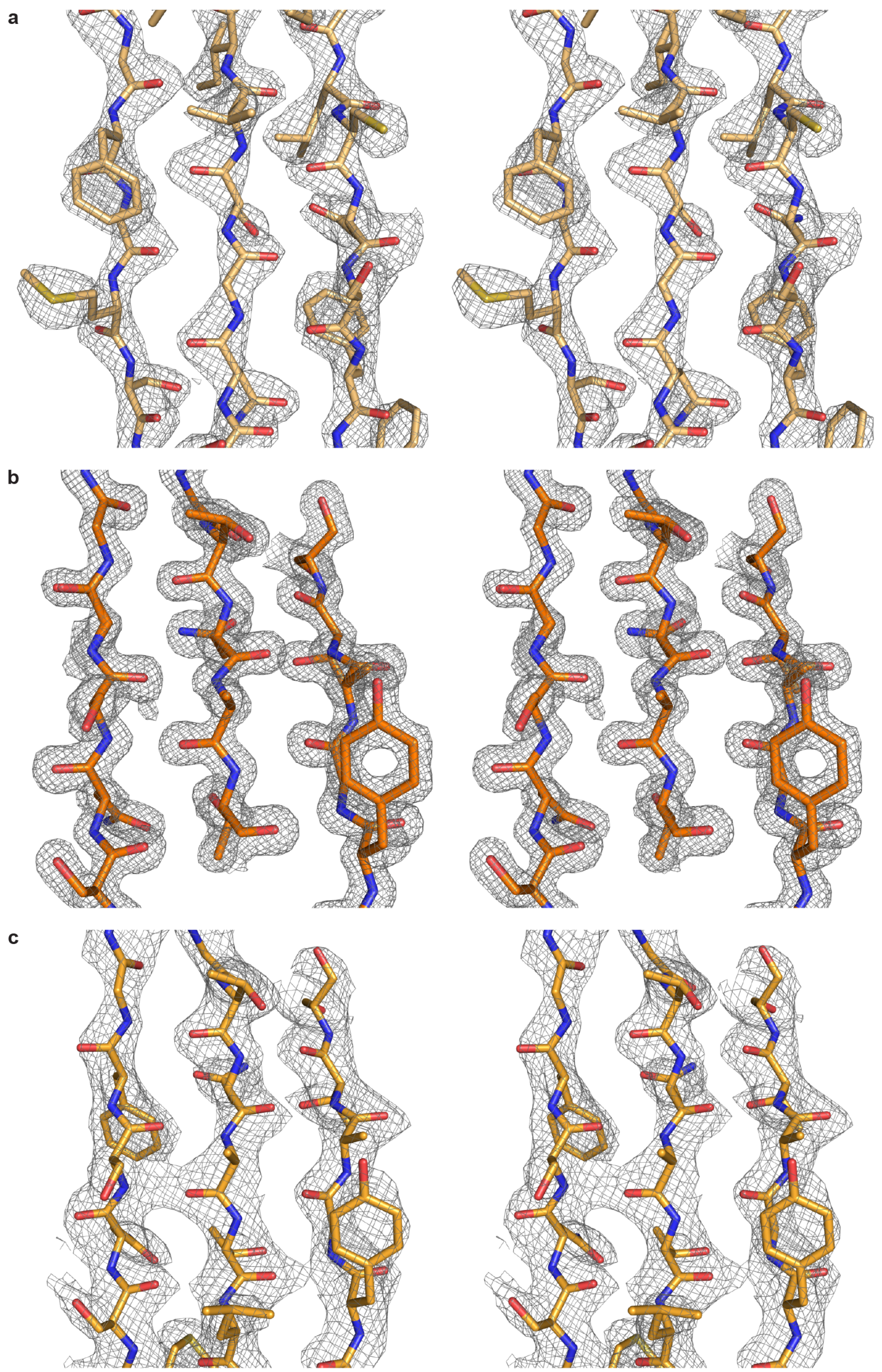


Electron density maps of reported structures. Stereo images of electron density maps for a. C16AcpP=FabF, (PDB: 6OKG, 2.3 Å). b. C12AcpP=FabB, (PDB: 6OKC ,1.55 Å) c. C16AcpP=FabB, (PDB: 6OKF, 2.5 Å.)
